# An orbitocortical-thalamic circuit suppresses binge alcohol-drinking

**DOI:** 10.1101/2024.07.03.601895

**Authors:** P Gimenez-Gomez, T Le, M Zinter, P M’Angale, V Duran-Laforet, TG Freels, R Pavchinskiy, S Molas, DP Schafer, AR Tapper, T Thomson, GE Martin

**Author notes:** Corresponding Authors: Gilles E. Martin. The Brudnick Neuropsychiatric Research Institute Department of Neurobiology University of Massachusetts Chan Medical School 364 Plantation street Worcester, MA 01605 Pablo Gimenez-Gomez. The Brudnick Neuropsychiatric Research Institute Department of Neurobiology University of Massachusetts Chan Medical School 364 Plantation street Worcester, MA 01605.

## Abstract

Alcohol consumption remains a significant global health challenge, causing millions of direct and indirect deaths annually. Intriguingly, recent work has highlighted the prefrontal cortex, a major brain area that regulates inhibitory control of behaviors, whose activity becomes dysregulated upon alcohol abuse. However, whether an endogenous mechanism exists within this brain area that limits alcohol consumption is unknown. Here we identify a discrete GABAergic neuronal ensemble in the medial orbitofrontal cortex (mOFC) that is selectively recruited during binge alcohol-drinking and intoxication. Upon alcohol intoxication, this neuronal ensemble suppresses binge drinking behavior. Optogenetically silencing of this population, or its ablation, results in uncontrolled binge alcohol consumption. We find that this neuronal ensemble is specific to alcohol and is not recruited by other rewarding substances. We further show, using brain-wide analysis, that this neuronal ensemble projects widely, and that its projections specifically to the mediodorsal thalamus are responsible for regulating binge alcohol drinking. Together, these results identify a brain circuit in the mOFC that serves to protect against binge drinking by halting alcohol intake. These results provide valuable insights into the complex nature of alcohol abuse and offers potential avenues for the development of mOFC neuronal ensemble-targeted interventions.

## Introduction

Alcohol remains the most widely consumed psychoactive drug worldwide, leading to 3 million deaths every year (5.3% of all deaths) and more than 200 associated diseases and injury conditions^1,2^. The last decades have advanced our knowledge about brain circuits and molecular mechanisms implicated in binge alcohol drinking^3^, a critical stage in the development of alcohol use disorders^4^. Unfortunately, the deeper knowledge of these processes has had a limited impact in the development of new treatments^5^. Current therapeutics that target entire neurotransmitter systems fail to work effectively, produce multiple side effects, and are unspecific since they regulate multiple brain areas^6^. Thus, there is a critical necessity to identify new, more specific molecular and cellular mechanisms underlying alcohol consumption and the development of alcohol abuse. Neuronal ensembles are discrete groups of coactive neurons evoked by a sensory stimuli or behaviors^7^ and constitute the basic units of neural code^8^. Although there is strong evidence that binge alcohol drinking activates discrete neuronal populations in several brain regions such as the medial orbitofrontal cortex (mOFC)^9^, the functional roles of those neurons remain unknown. The prefrontal cortex, a structure that enables decision making, executive function and inhibitory control^10^, is one of the most vulnerable brain regions to the effects of alcohol^11^. The general view is that binge alcohol drinking impairs mOFC circuits affecting cognitive functions^12^. However, due to the key role of this area in executive functions and inhibitory control, it is possible that during early stages of drinking, mOFC may activate circuits at a neuronal ensemble level in response to alcohol that could limit its consumption. Harnessing such circuits could be the basis of a new therapeutic approach. To explore this possibility, here we aimed to identify neuronal ensembles that control binge alcohol drinking in the mOFC.

## Results

### Binge alcohol drinking recruits a stable neuronal ensemble in the mOFC

Although neuronal activity in the mOFC has been associated with binge alcohol drinking in humans^13^ and preclinical models^14,15^, the exact mechanism on how neuronal ensembles in the mOFC mediates the effects of binge drinking remain elusive. To study the role of the mOFC in binge alcohol-drinking, C57BL/6J mice (8 weeks old) were subjected to the *Drinking in the Dark* (DID) paradigm^16^ (Extended Data Fig. 1a). Mice had access to alcohol containing drinking solutions for 2 h over a 3-day period, followed by a 4 h alcohol exposure on day 4 that resulted in high levels of alcohol consumption. As a control, separate groups of animals were exposed to saccharine (sacch) or water drinking solutions for the same period. We used sacch as a control since it is a rewarding substance that do not produce addiction. All groups of mice increased liquid consumption on the last DID session, with no differences between males and females (Extended Data Fig. 1b-d). Importantly at the end of the 4 h session, blood ethanol concentration (BEC) reached concentrations higher than 80 mg/dl (Extended Data Fig. 1d), levels defined as intoxicating levels^17^ in humans during an episode of binge alcohol drinking, validating this model to study the formation of neuronal ensembles during binge alcohol drinking. To investigate the response of mOFC to binge alcohol drinking, we performed a time-course expression analysis of the Fos neuronal activation marker on the last day of consumption (Extended Data Fig. 1e). We found that both alcohol and sacch activated a discrete Fos^+^ neuronal ensemble in the mOFC 2 h after consumption, but not at earlier or later time points (Extended Data Fig. 1f and 1g), with no differences between males and females (Extended Data Fig. 1h). In the water control group, the activated neuronal ensemble was nearly undetectable (Extended Data Fig. 1f and 1g). The mOFC neuronal ensemble activated by sacch or binge alcohol drinking comprised 42 ± 5 and 68 ± 10 neurons per field of view, respectively (Extended Data Fig. 1i). Considering there were no differences between males and females in the amount of alcohol intake when normalized per weight and the size of the neuronal ensemble, both sexes were included in the rest of the study.

To study the functional role of alcohol neuronal ensemble in the mOFC, we used the TRAP2 mouse model^18^. This model enables the transient activation of the cre recombinase driven by the Fos locus allowing permanent genetic access to neurons activated by binge alcohol drinking. When crossed with the Ai14 transgenic mouse line, it allows permanent expression of the tdTomato reporter upon neuronal activation and 4-hydroxitamoxifen (4-OHT) injection (Fig. 1a). To induce cre expression, we injected 4-OHT 2 h after the last day of binge alcohol drinking and examined tdTomato expression 10 days later to allow for its robust expression (here referred as TRAPed neurons) (Fig. 1b). For comparisons, we included the water and sacch control groups. TRAP2 mice consumed 47.35 ± 7.9 ml/kg of water (Fig. 1c) 66.75 ± 13.73 ml/kg of sacch (Fig. 1d) and 6.73 ± 0.89 g/kg of alcohol on the last day of the DID protocol and reached a BEC of 98.63 ± 11.46 mg/dl indicating intoxication levels (Fig. 1e). Mice drank similar amounts as we described in C57BL/6J mice, suggesting that the transgene expression did not disrupt the pattern of binge drinking behavior. In the mOFC of TRAP2 mice, the number of TRAPed neurons represented 2% ± 0.4 and 3.78% ± 0.27 neurons of the total DAPI^+^ cells population for sacch and binge alcohol drinking respectively (Fig. 1f-g).

**Fig. 1.**
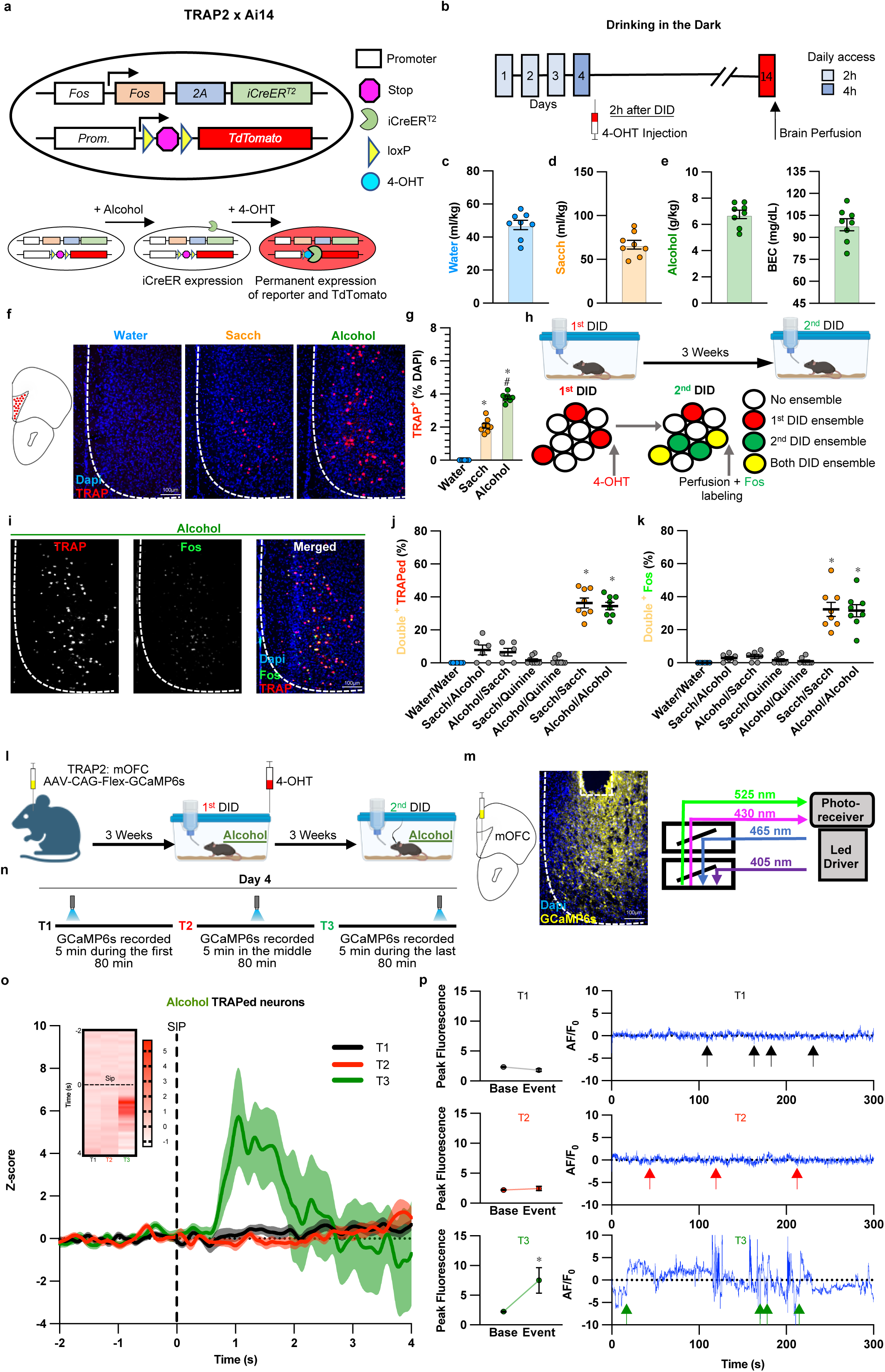
Binge alcohol drinking recruits a stable neuronal ensemble in the mOFC. **a.** Schematic of the strategy to TRAP the neurons activated in response to binge alcohol drinking. **b.** Timeline for TRAPping Fos^+^ neurons. **c.** Water consumption at the end of the last day of DID in control mice (n=8). **d.** Sacch consumption at the end of the last day of DID in control mice (n=8). **e.** Alcohol consumption and BEC at the end of the last day of DID (n=8,8). **f.** Representative images of TRAPed neurons expression after water, sacch and alcohol consumption in the mOFC. The broken white lines show the medial boundary of the mOFC. **g.** TRAPed neurons (expressed as a % of the total DAPI cells) for water, sacch and alcohol groups (*F*_(20,22)_ = 339.4, *p* < 0.0001; * *vs* Water Group, # *vs* Sacch Group; *n*=8,8,8). **h.** Schematic of the strategy to study neuronal ensemble stability. We TRAPed neurons at the end of the first drinking episode (1^st^ DID) following 4-OHT injection. Three weeks later, we assessed reactivation at the end of second drinking episode (2^nd^ DID) by measuring Fos immunofluorescence. **i.** Representative images from the Alcohol/Alcohol group of TRAPed neurons (in gray), Fos^+^ neurons (in gray) and TRAPed and Fos^+^ neurons overlap. The broken white lines show the medial boundary of the mOFC. **j.** Quantification of percentage of TRAPed neurons that are Fos+ neurons (*F*_(6,44)_ = 67.43, *p* < 0.0001; * *vs* Water/Water group, *n*=8,6,6,8,8,8,8). **k.** Quantification of percentage of Fos^+^ neurons that are TRAPed neurons (*F*_(6,43)_ = 36.08, *p* < 0.0001; * *vs* Water/Water group, *n*=8,6,6,8,8,8,8). **l.** Timeline of fiber photometry experiments in TRAPed neurons after binge alcohol drinking consumption. **m.** Representative image of the expression of GCaMP6s-eGFP. The broken white lines show the medial boundary of the mOFC and the optic fiber trace. Schematic of the experimental approach used to measure GCaMP6s signal as indicative of TRAPed neurons activity during drinking. **n.** GCaMP6s fluorescence was recorded at three different time points: T1 (5 min randomly selected during the first 80 min), T2 (5 min randomly selected in the middle 80 min) or T3 (5 min randomly selected during the last 80 min). **o.** Mean mOFC GCaMP6s-recorded calcium activity in TRAPed neurons 2 seconds prior to the sip (vertical broken line) and 4 seconds after the sip in T1, T2 and T3. Insert shows heatmap raster plot of mOFC GCaMP6s-recorded calcium activity in TRAPed neurons in a representative mouse at T1, T2 and T3. **p.** Peak Fluorescence during base (−2 to 0 seconds) and event (0 to 4 seconds) (T1: *t*_(94)_= 1.9, *p*=0.06 and T2: *t*_(84)_= 1.4, *p*=0.16 *ns*; T3: *t*_(61)_= 2.345, *p*=0.0223; * *vs* Base; *n*=12,12,12). Representative recorded GCaMP6s activity in the mOFC during 5 min in T1, T2 and T3. Colored arrows represent a sip. In all plots and statistical tests, *n* represents biologically independent animals (except for **o** and **p** that represent the average of animals), *n* in caption represents biologically independent animals. Summary graphs show mean ± s.e.m.

Neuronal ensembles consist of sparse, distinct populations of neurons active during a particular experience and which reactivation drives behavior^19^. We explored whether TRAPed neurons recruited during the first binge drinking episode (1^st^ DID, Fig. 1h) could be reactivated in mice exposed to another binge drinking episode (2^nd^ DID, Fig. 1h). To test this idea, we injected 4-OHT at the end of the 1^st^ DID to TRAP the binge alcohol drinking neuronal ensemble and performed immunofluorescence staining at the end of the 2^nd^ DID to label Fos+ neurons. This strategy allowed us to distinguish neurons exclusively activated during the 1^st^ DID (TRAPed, red), versus those reactivated during the 2^nd^ DID episode (TRAPed/Fos^+^, yellow), or solely activated during the 2^nd^ DID (Fos^+^, green, Fig. 1h). Exposure to a 2^nd^ DID episode significantly reactivated neuronal ensembles both in the binge alcohol drinking and the sacch groups, measured as a fraction of total TRAPed neurons (Fig. 1i-j) or Fos+ neurons (Fig. 1k) indicating that the neuronal ensemble is stable among different DID episodes. Because the level of TRAPed neurons in water-drinking mice was undetectable we used the sacch neuronal ensemble as a control for the rest of the study. Importantly, the alcohol TRAPed neurons did not reactivate upon exposure to sacch or quinine, rewarding and aversive drinking solutions respectively. Finally, we found no overlap between alcohol and sacch neuronal ensembles indicating that alcohol and sacch activate independent neuronal ensembles (Alcohol/Sacch, Fig. 1i,k). Taken together those data demonstrates that the alcohol neuronal ensemble is stable among different DID episodes.

Alcohol has been associated with several high-risk behaviors^20^. To determine if the alcohol neuronal ensemble could be reactivated by some of those behaviors, three weeks following binge alcohol drinking, we exposed mice to stress, nicotine consumption, social defeat (SD) or social exploration (behaviors associated with and/or impacted by alcohol) (Extended Data Fig. 2a). The alcohol neuronal ensemble was partially reactivated upon a subsequent binge alcohol drinking episode when measured as a fraction of total TRAPed neurons or Fos^+^ neurons (Extended Data Fig. 2b-d). However, none of the other conditions reactivated the alcohol neuronal ensemble (Extended Data Fig. 2b-d). We wondered whether these conditions, when presented before mice were allowed to binge drinking alcohol, could change the size of the alcohol neuronal ensemble (Extended Data Fig. 2e). We found no differences in the size of the neuronal ensemble (Extended Data Fig. 2f and 2g) and the alcohol consumed (Extended Data Fig. 2h). Altogether these results indicate that the alcohol neuronal ensemble is exclusive for alcohol, and it is not affected by related behaviors.

### Alcohol intoxication is required for the mOFC neuronal ensemble activation

As the first set of experiments to identify the alcohol ensemble was done by static imaging of activity-dependent of immediate early genes, we used fiber photometry to determine when TRAPed neurons were activated during the 2^nd^ DID, and their precise temporal relationship to drinking (referred here as SIP). We injected a viral vector into the mOFC containing a plasmid coding for the calcium indicator GCaMP6s and induced its selective expression in TRAPed neurons following 4-OHT injection at the end of the 1^st^ DID. Three weeks later, mice were allowed to drink alcohol while calcium signals were recorded via a cannula implanted above the mOFC during the last session of a 2^nd^ DID episode (Fig. 1l-m). We divided the 4 h drinking session in three periods: T1 (0 to 80 min), T2 (80 to 160 min) or T3 (160 to 240). For each period we recorded calcium signals for 5 min and we time-locked activity of the alcohol TRAPed neurons to drinking (Fig. 1n). Alcohol TRAPed neurons significantly increased activity during T3 with no changes at earlier time points (Fig. 1o-p). Interestingly, no changes in calcium dynamics in alcohol TRAPed neurons were detected if animals were presented to water, sacch or quinine during T3 on the 2^nd^ DID episode (Extended Data Fig. 3a-e). Considering that TRAPed neurons calcium response sharply increased only towards the end of the alcohol drinking episode, we wondered whether intoxication level was necessary for those neurons to show activity. To explore this possibility, we compared alcohol drinking patterns and the formation of mOFC alcohol neuronal ensembles in mice allowed to drink 10% or 20% alcohol solutions in the DID paradigm (Extended Data Fig. 4a). Mice receiving 10% alcohol consumed less than those that drank 10% alcohol (Extended Data Fig. 4b). Moreover, only mice that drank 20% alcohol reached intoxication BEC levels (Extended Data Fig. 4c). Remarkably, in mice exposed to the 10% alcohol solution, we did not observe the formation of the alcohol neuronal ensemble in the mOFC (Extended Data Fig. 4d-e). To validate that the formation of the alcohol neuronal ensemble depends on alcohol intoxication, we injected Pyrazole, a compound that blocks the metabolism of alcohol^21^, in mice that had access to 20% alcohol for only 2 h (Extended Data Fig. 4f). Although these mice drank less alcohol than those that had 4 h access (Extended Data Fig. 4g), their BEC reached the same alcohol intoxication levels (Extended Data Fig. 4h). Thus, mice that drank alcohol for 2 h formed the alcohol neuronal ensemble only when they also received pyrazole (Extended Data Fig. 4i-j), demonstrating that a high BEC is necessary for the neuronal ensemble formation. In contrast, sacch neuronal ensemble was not sensitive to the amount of sacch consumed (Extended Data Fig. 4k-n). These results indicate that binge alcohol drinking recruits a stable neuronal ensemble in the mOFC which formation and reactivation depend on the levels of alcohol intake.

### The alcohol ensemble neurons are predominantly GABAergic neurons highly permeable to calcium

As the mOFC is a molecularly heterogenous brain region^22,23^ we used single-cell RNA seq (scRNA-seq) to examine transcripts expressed in TRAPed and non-TRAPed neurons of alcohol and sacch drinking mice. 10 days after binge drinking, we micro-dissected the mOFC and enzymatically digested the tissue to obtain well-dissociated, highly viable single cells (viability > 70%). We captured single-cell transcriptomes and generated cDNA libraries for posterior sequencing (Fig. 2a). We recovered the transcriptomes from a total of 38960 cells across 4 independent samples. Principal-components analysis was used to reduce dimensionality of the data, followed by graph-based clustering, and visualization with the UMAP algorithm (Fig. 2b). This method identified 15 cell clusters based on their canonical gene markers (Fig. 2c). As a strategy to identify the TRAPed neurons, we used the expression of the *“Flox-stop”* sequence. We identified 643 TRAPed and 4260 non TRAPed neurons across 7 neuronal clusters (Fig. 2d). Regarding the alcohol group, we found that most alcohol TRAPed neurons showed substantial expression of the transcript encoding for GAD2 (Fig. 2e), a result that was validated by immunostaining with the GAD2 marker indicating that the alcohol neuronal ensemble is mostly formed by GABAergic cells (Fig. 2f).

**Fig. 2.**
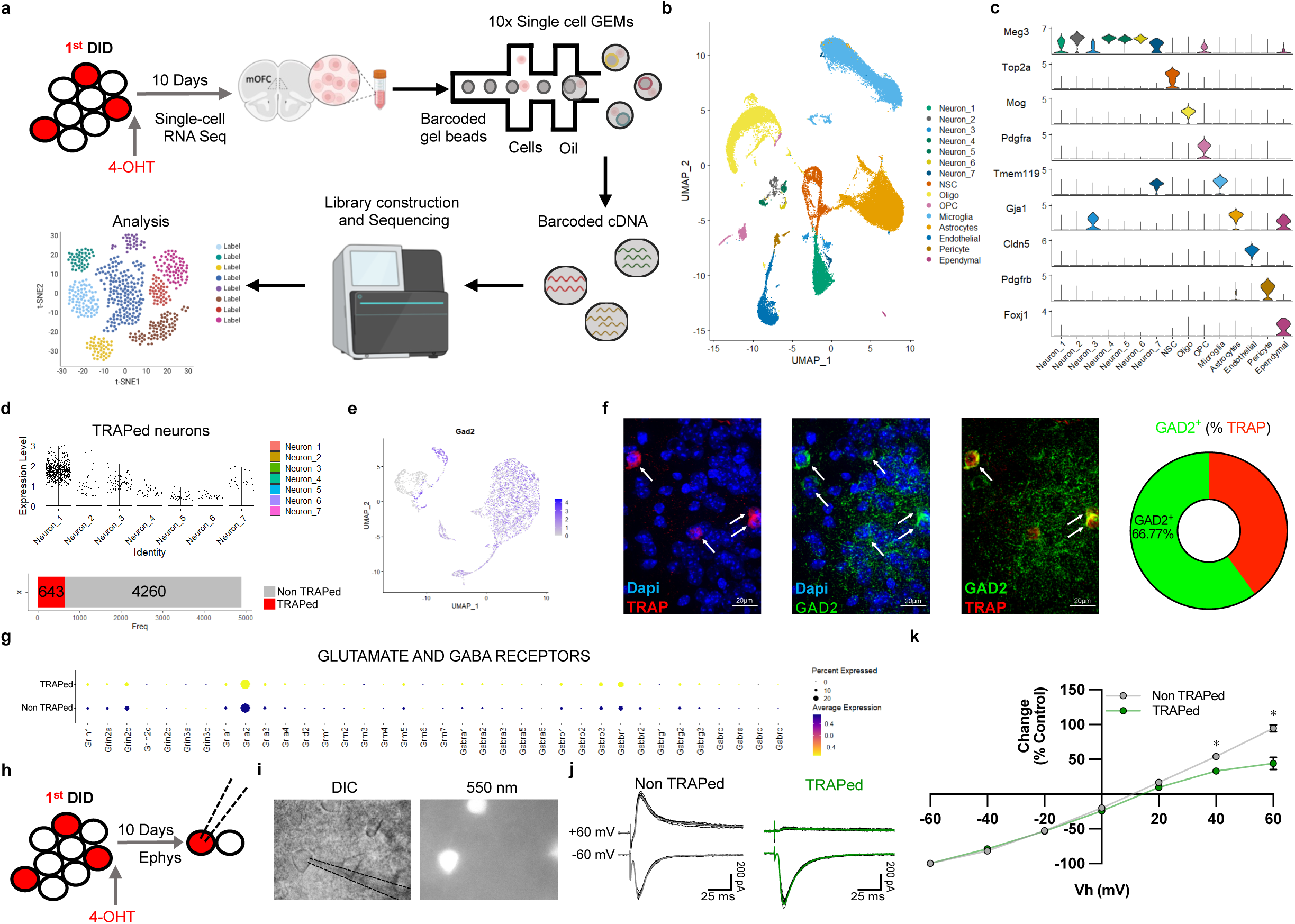
The alcohol neuronal ensemble is composed of GABAergic neurons highly permeable to calcium. **a.** Schematic of scRNA-seq experiments. **b.** UMAP dimensional reduction and visualization of transcriptional profiles of mOFC cells (18430 and 20530 cells from binge alcohol drinking mice and sacch drinking mice, respectively; *n*=4). **c.** Violin plots showing levels of expression of canonical marker genes in neuronal and non-neuronal cells. **d.** Strategy to identify the TRAPed neurons across neuronal clusters using the Floxed-Stop sequence and number of TRAPed neurons identified. **e.** Feature plot showing normalized expression values for GAD2 across the TRAPed neurons in the UMAP space. **f.** Representative images of TRAPed neurons, GAD2^+^ neurons, overlap TRAPed/GAD2^+^ neurons and proportion of GAD2^+^ neurons in the alcohol neuronal ensemble. **g.** Dot plot of scaled expression levels (color) and the proportions of expressing neurons (dot size) of glutamate and GABA-receptor-related genes. **h.** Schematic of electrophysiology experiments. **i.** Representative images of mOFC neurons in DIC (left panel) and excited with green light (550 nm; right panel). **j.** Representative AMPA-EPSCs from non-TRAPed and TRAPed neurons recorded in the presence of NMDA and GABA_A_ receptor antagonists at holding potentials of -60 and +60mV. EPSCs were evoked every 10 sec 8 consecutive times. Average traces are shown in gray and green for non TRAPed and TRAPed neurons, respectively. **k.** Current-voltage relationship of AMPA current between -60 and +60 mV. Values are expressed as percent of current evoked at -60 mV. (*F*_group(1,144)_ = 29.52, *p* < 0.0001; *F*_Vh(3,80)_ = 533.1, *p* < 0.0001; *F*_interaction(6,144)_ =11.78, *p* < 0.0001; * *vs* non TRAPed same time; *n*=12,12). In all plots and statistical tests *n* represent cells (except for **k** that represent the average of cells), *n* in caption represents biologically independent animals. Summary graphs show mean ± s.e.m.

We next sough to better understand why some neurons showed activity and constituted a neuronal ensemble while others remained inactive after binge alcohol drinking. To do so, we measured the transmission of the two most prevalent neurotransmitters in the prefrontal cortex, glutamate and GABA, in alcohol TRAPed and non-TRAPed neurons. We first compared the expression of transcripts coding for glutamate and GABA receptor subunits. Expression for GRIA2, the transcript that encodes the AMPAR receptor subunit that regulates the channel calcium permeability, was downregulated in TRAPed neurons compared to non-TRAPed neurons (Fig. 2g), suggesting that AMPAR permeability to calcium in TRAPed neurons was higher than that of non-TRAPed neurons. We validated this result by recording AMPA currents at the same time point as the scRNA-seq experiment (Fig. 2h-j). While AMPA current-voltage relationship was linear in non-TRAPed neurons, TRAPed neurons displayed a strong rectification, supporting the notion that AMPARs in TRAPed neurons are more permeable to calcium than their non-TRAPed counterparts (Fig. 2k). Interestingly, using the same approach (Extended Data Fig. 5a) we found no significant differences in the NMDA current-voltage relationship (Extended Data Fig. 5b) and the AMPA/NMDA ratio (Extended Data Fig. 5c) between TRAPed and non-TRAPed neurons. Finally, gene expression of sodium and potassium channel subunits revealed a downregulation of the transcripts coding for the sodium channel subunit Scn3b in TRAPed neurons vs non TRAPed neurons (Extended Data Fig. 5d). Electrophysiological recordings of action potentials showing a significant reduction of their amplitude with no alteration in other parameters supported this result (Extended Data Fig. 5e-n). Interestingly, sacch TRAPed neurons, as in the alcohol neuronal ensemble, are GABAergic neurons (Extended Data Fig. 6a-b) and show the same permeability for calcium (Extended Data Fig. 6c-g) with no alteration in the NMDA current (Extended Data Fig. 6h) although their A/N rate was significantly smaller compared to non-TRAPed neurons (Extended Data Fig. 6i). Finally, we found a similar pattern of expression of the genes related to ion channels (Extended Data Fig. 6j) and in the passive and active membrane properties (Extended Data Fig. 6k-t). These results showed that the binge alcohol drinking and sacch neuronal ensembles are similar as both are formed mostly by GABAergic neurons that present alterations in their active membrane properties and a decrease in the expression of the GluA2 subunit, a subunit controlling AMPA receptor calcium permeability which may be critical for the formation of the neuronal ensembles.

### mOFC alcohol neuronal ensemble suppresses alcohol consumption to control binge alcohol drinking

To test the functional role of the alcohol neuronal ensemble in binge alcohol drinking, we expressed ChR2 in TRAPed neurons and optogenetically stimulated these neurons during the last 4 h session of a 2^nd^ DID episode (Fig. 3a-b). Photostimulation occurred in a closed-loop manner which turned ON each time a mouse contacted the spout delivering alcohol and remained ON for 2 s after the lick (Fig. 3c). Again, we divided the last 4 h drinking session in 3 periods (Fig. 3d). Photostimulation of TRAPed neurons significantly decreased alcohol intake and the BEC at all time points as compared to control mice expressing eYFP (Fig. 3e), indicating that the role of these neurons is to suppress binge alcohol drinking. Although photostimulation of TRAPed neurons did not alter locomotor activity of mice (Fig. 3f), a real-time place preference assay showed that mice displayed avoidance in the photostimulated-paired side of the chamber when compared with control mice expressing eYFP, indicating that the activation of those neurons is aversive (Fig. 3g). In addition, photostimulation of the alcohol neuronal ensemble reduced the preference for alcohol in a choice paradigm, when animals were assessed for voluntary alcohol consumption (Extended Data Fig. 7a-b). Intake of water, quinine, and sacch were not affected by alcohol neuronal ensemble photostimulation (Extended Data Fig. 7c-e). On the other hand, photostimulation of the TRAPed neurons associated with sacch consumption did not alter sacch intake in a 2^nd^ DID episode (Fig 3h), neither it affected water, quinine or alcohol consumption (Extended Data Fig. 7f-j), locomotor activity (Extended Data Fig. 7u) or induced changes in the real time place preference test (Extended Data Fig. 7v). To test the necessity of TRAPed neurons on binge alcohol drinking, we expressed NpHR in TRAPed neurons and silenced their activity in a closed-loop manner paired with alcohol drinking events (sip) (Fig. 3i-l). Photoinhibition of the TRAPed neurons significantly increased alcohol intake and the associated BEC in all the time points measured (Fig. 3m), with no alteration of locomotor activity (Fig. 3n). In a real-time place preference assay, mice in which the neuronal ensemble was silenced, displayed a stronger preference for the photoinhibited-paired side of the chamber compared to control mice expressing eYFP demonstrating that the inhibition of this selective mOFC neuronal ensemble induces rewarding effects (Fig. 3o). Photoinhibition of TRAPed neurons also increased the preference for alcohol when animals where offered a choice (Extended Data Fig. 7k-l) but did not affect the consumption of other substances (Extended Data Fig. 7m-o). In contrast, photoinhibition of the mOFC sacch neuronal ensemble did not alter total sacch intake in a 2^nd^ DID (Fig. 3p) nor impacted the consumption of other substances (Extended Data Fig. 7p-t), locomotor activity (Extended Data Fig. 7w) or real-time place preference (Extended Data Fig. 7x), further validating the idea that the sacch neuronal ensemble does not regulate sacch consumption. These results demonstrate that activity of the alcohol neuronal ensemble is necessary to suppress binge drinking and establish the role of this neuronal ensemble as a suppressor of binge alcohol drinking.

**Fig. 3.**
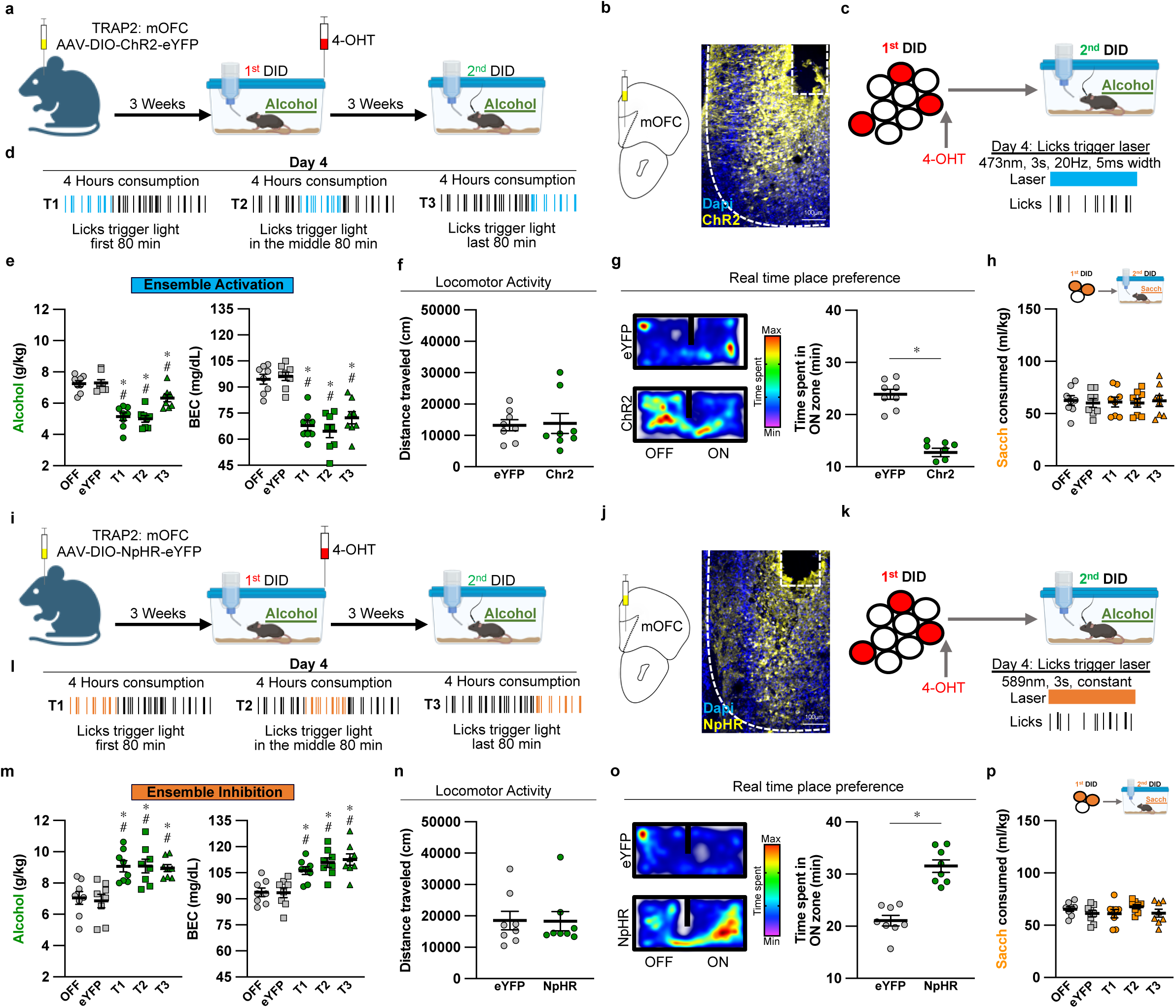
The alcohol neuronal ensemble suppresses binge alcohol drinking. **a.** Timeline of optogenetics experiments in ChR2-expressing TRAPed neurons during binge alcohol consumption. **b.** Representative image of the expression of ChR2-eYFP in the mOFC. The broken white lines show the medial boundary of the mOFC and the optic fiber trace. **c**. Pattern of light delivered to activate TRAPed neurons during the last day of the second DID. Blue light stimulation (473 nm, 20 Hz and 5 ms width) is triggered when mice establish contact with alcohol bottle spouts. **d.** TRAPed neurons were optogenetically stimulated at three different time points: T1 (first 80 min), T2 (middle 80 min) or T3 (last 80 min). **e.** Alcohol consumption (*F*_(4,35)_ = 25.67, *p* < 0.0001; * *vs* OFF group, # *vs* eYFP group *n*=8,7,8,8,8) and BEC measured the last day of the second DID while the mice were photostimulated (*F*_(4,35)_ = 23.25, *p* < 0.0001; * *vs* OFF group, # *vs* eYFP group *n*=8,8,8,8,8). **f.** Locomotor activity of mice during photostimulation (*t*_(14)_= 0.15, *p*=0.87; *n*=8,8). **g.** Real-time preference/avoidance test of mice during photostimulation (*t*_(14)_= 8.93, *p*<0.0001; *n*=8,8) and representative images. **h.** Photostimulation of the sacch TRAPed neurons during a second episode of sacch consumption (*F*_(4,35)_ = 0.057, *p* = 0.99; *n*=8,8,8,8,8). **i.** Timeline of optogenetics experiments using NpHR in TRAPed neurons during binge alcohol drinking consumption. **j.** Representative image of the expression of NpHR-eYFP in the mOFC. The broken white lines show the medial boundary of the mOFC and the optic fiber trace. **k**. Pattern of light delivered to inhibit TRAPed neurons during the last day of the second DID. Red light stimulation (589 nm, 3s, constant) is triggered when mice establish contact with alcohol bottle spouts. **l.** TRAPed neurons were optogenetically inhibited at three different time points: T1 (first 80 min), T2 (middle 80 min) or T3 (last 80 min). **m.** Alcohol consumption (*F*_(4,35)_ = 9.072, *p* < 0.0001; * *vs* OFF group, # *vs* eYFP group *n*=8,8,8,8,8) and BEC measured the last day of the second DID while the mice were photoinhibited (*F*_(4,35)_ = 11.52, *p* < 0.0001; * *vs* OFF group, # *vs* eYFP group *n*=8,8,8,8,8). **n.** Locomotor activity of mice during photoinhibition (*t*_(14)_= 0.04, *p*=0.96; *n*=8,8). **o.** Real-time preference/avoidance test of mice during photoinhibition (*t*_(14)_= 6.7, *p*<0.0001; *n*=8,8) and representative images. **p.** Photoinhibition of the sacch TRAPed neurons during a second episode of sacch consumption (*F*_(4,35)_ = 0.94, *p* = 0.444; *n*=8,8,8,8,8). In all plots and statistical tests *n* represent biologically independent animals, *n* in caption represents biologically independent animals. Summary graphs show mean ± s.e.m.

Because the optogenetic approach demonstrated that the alcohol neuronal ensemble controls alcohol intake, we wanted to know whether it could also affect long-term consumption. To address this question, we ablated the neuronal ensemble using a cre-dependent casp3 approach (Extended Data Fig. 8a-b). We compared the consumption in the 1^st^ DID (formation of the neuronal ensemble) and 2^nd^ DID after ablation of the neuronal ensemble by casp3 (Extended Data Fig. 8c). This approach sharply decreased the size of the neuronal ensembles measured in both alcohol and sacch drinking mice, confirming the validity of this approach to study long-term consumption (Extended Data Fig. 8d-e). Similar to our optogenetic experiments, ablation of the alcohol neuronal ensemble increased alcohol consumption in a 2^nd^ DID episode (Extended Data Fig. 8f). This contrast with the ablation of the sacch neuronal ensemble that failed to alter sacch consumption (Extended Data Fig. 8g). Ablation of the alcohol neuronal ensemble did not alter water consumption (Extended Data Fig. 8h) or locomotor activity (Extended Data Fig. 8i) but did significantly increase the preference for alcohol in a choice paradigm (Extended Data Fig. 8j). We then investigated whether mice with ablated alcohol neuronal ensemble would maintain their alcohol drinking despite negative consequences by adding increasing doses of quinine to the alcohol solution^24^. Mice with alcohol neuronal ensemble ablation showed a robust resistance to quinine adulteration indicating that the alcohol neuronal ensemble is necessary to maintain the negative valence of alcohol consumption (Extended Data Fig. 8k). Finally, we measured long term consumption demonstrating a robust increase in long term consumption after the ablation of the neuronal ensemble (Extended Data Fig. 8l). Taken together, these data demonstrate that ablation of the alcohol neuronal ensemble has a stable, long-lasting effect on alcohol consumption in subsequent DID episodes.

Our transcriptomic and electrophysiological data suggested the alcohol neuronal ensemble was mostly comprised of GABAergic neurons. To test whether the mOFC controls alcohol consumption via a selective alcohol neuronal ensemble or through the overall mOFC GABAergic interneuron population, we performed optogenetic experiments in GAD2 mice using a similar approach as the described above (Extended Data Fig. 9a-b). Photostimulation using ChR2 of overall mOFC GABAergic neurons did not alter alcohol consumption (Extended Data Fig. 9c), locomotor activity (Extended Data Fig. 9d) or the preference for a chamber in a real-time place preference test (Extended Data Fig. 9e). Similarly, NpHR-mediated photoinhibition of the mOFC GABAergic neuronal population did not alter any of those behaviors (Extended Data Fig. 9f-j) further validating the idea that modulation of alcohol consumption is caused by the alcohol neuronal ensemble and not for all the GABAergic neurons in the mOFC. Those data shows that the alcohol neuronal ensemble is related to binge alcohol drinking and that this neuronal ensemble is necessary and sufficient to suppress alcohol consumption.

### The alcohol neuronal ensemble projects preferentially to PAG and MD

Neurons in the mOFC are highly diverse in their projection-specific pattern and functional connectivity that may reflect their complex role in cognitive flexibility, inhibitory control, and decision-making processes^25^. To determine where they projected, we injected in the left mOFC AAVs containing the hSyn.Flex.mGFP.2A.synaptophysin.mRuby construct. Following cre recombination, synaptophysin fused to the mRuby red fluorophore was selectively transported into the axonal compartments of the neuronal ensemble^26^. After an initial episode of alcohol or sacch consumption we induced cre expression in TRAPed neurons by injecting 4-OHT. 4 weeks later we used iDISCO+ to clear the brain and label mRuby (Fig. 4a-b)^27^. We then paired lightsheet microscopy with SmartAnalytics software analysis^28^ to quantify the brain-wide axonal projections of mOFC alcohol and sacch neuronal ensembles (Fig. 4c). In sharp contrast to the sacch neuronal ensemble that only projected to a few areas (Fig. 4d-f), alcohol TRAPed neurons projected broadly (Fig. 4d) with dense innervation in periaqueductal gray area (PAG), visual areas and thalamic nuclei (Fig 4e-f and Tables s1-2). PAG received the most innervation from the alcohol neuronal ensemble, followed by the mediodorsal thalamus (MD). Interestingly, while projections to the PAG in alcohol and sacch drinking mice were similar, projections to the MD in alcohol mice was significantly higher compared to those in sacch mice. Therefore, we used IF to validate projections specifically to these regions with higher resolution than whole-brain mapping. We found that projections to the PAG from mOFC binge alcohol drinking and sacch neuronal ensembles were similar (Fig. 4g), whereas the MD received significantly more innervations from the mOFC alcohol neuronal ensemble as compared to the sacch neuronal ensemble (Fig. 4h). Taken together our results reflects a differential pattern of projections that could explain the effect of mOFC alcohol TRAPed neurons suppressing alcohol consumption.

**Fig. 4.**
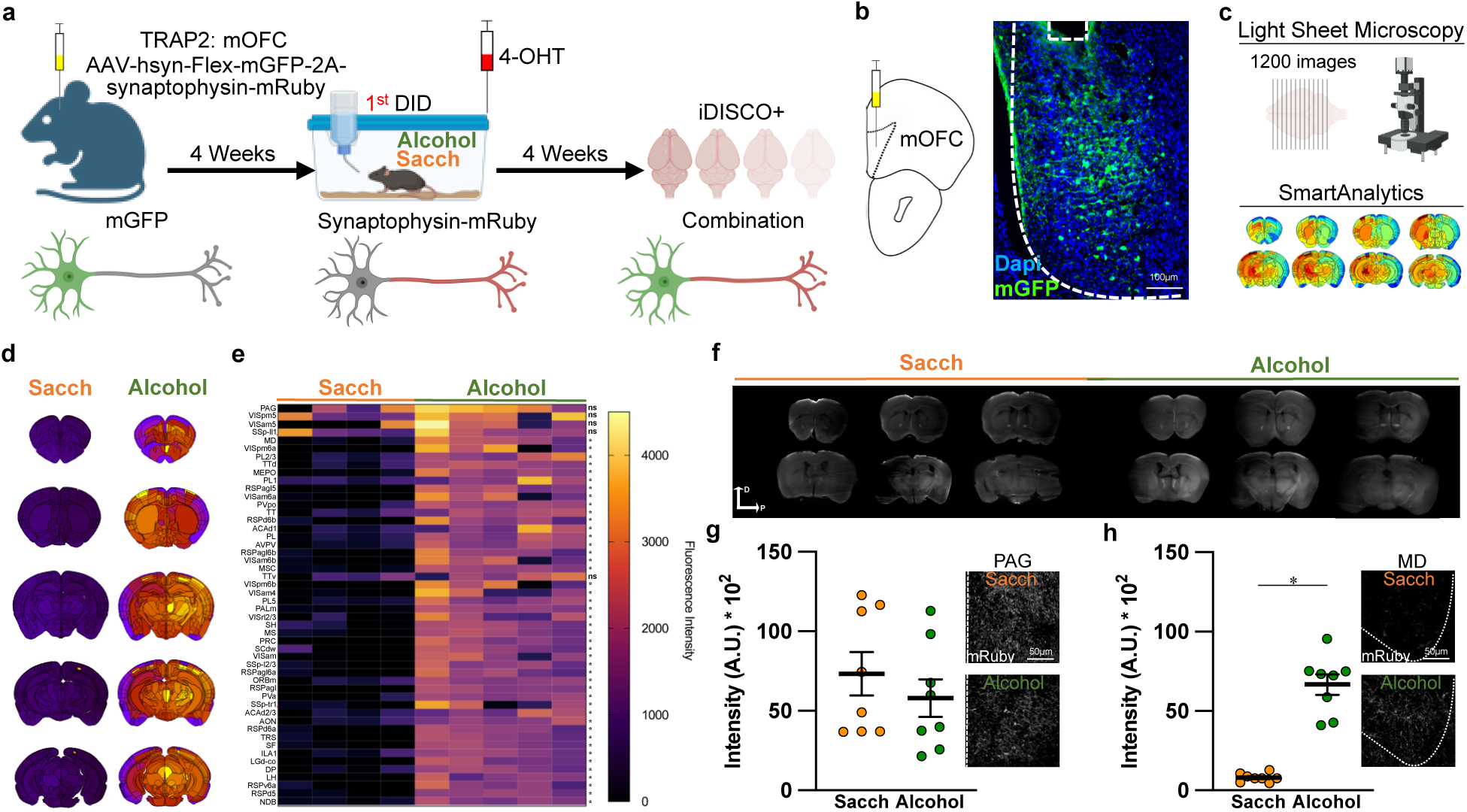
The alcohol neuronal ensemble projects preferentially to PAG and MD. **a.** Timeline for iDISCO+ experiments. **b.** Representative image of the expression of AAV-hsyn-Flex-mGFP-2A-synaptophysin-mRuby in the mOFC. The broken white lines show the medial boundary of the mOFC and the injection site. **c.** We combined Light Sheet Microscopy and Smart Analytics (Life Canvas) to quantify the complete projectome of the alcohol neuronal ensemble. **d**. Representative brains obtained from sacch and alcohol groups using Smart Analytics. **e.** Heatmap showing axonal innervation of TRAPed neurons after sacch or alcohol consumption (n=4,5; * p< 0,05). **f.** Representative images of brains obtained using Light Sheet microscopy. **g.** Quantification of signal intensity in arbitrary units (indicative of axonal innervation) in PAG (*t*_(14)_= 0.84, *p*=0.41; *n*=8,8). **h.** Quantification of signal intensity in arbitrary units (indicative of axonal innervation) in MD (*t*_(14)_= 8.94, *p*<0.0001; *n*=8,8). In all plots and statistical tests, *n* represents biologically independent animals. Summary graphs show mean ± s.e.m.

### mOFC neuronal ensemble projections to the mediodorsal thalamus suppress alcohol consumption

After identifying the projections of mOFC alcohol neuronal ensemble, we next sough to test the functional relevance of these projections to PAG and MD, two regions that have been implicated in alcohol consumption^29,30^. First, we investigated the role of mOFC→PAG projections by expressing ChR2 in the mOFC alcohol TRAPed neurons and optogenetically stimulated their axon terminals in the PAG each time the mice contacted the spout delivering alcohol during a 2^nd^ DID episode (Extended Data Fig. 10a-b). We delivered a pulse of light to photostimulate the axons of mOFC alcohol TRAPed neurons in PAG during T1 (Extended Data Fig. 10c). Photostimulation of the mOFC alcohol TRAPed neurons projecting to the PAG did not alter alcohol consumption, BEC (Extended Data Fig. 10d), locomotor activity (Extended Data Fig. 10e) or real-time place preference test (Extended Data Fig. 10f). Similarly, photoinhibition of the mOFC→PAG neuronal ensemble projections failed to alter alcohol consumption (Extended Data Fig. 10g-l). Next, we examined the role of mOFC alcohol TRAPed neurons projections to the MD, an area that receives afferents from the alcohol neuronal ensemble but not from the sacch neuronal ensemble. We expressed ChR2 in the mOFC alcohol TRAPed neurons and optogenetically stimulated the projections to the MD each time the mice contacted the spout delivering alcohol during a 2^nd^ DID episode of binge alcohol drinking (Fig. 5a-b). We found that the photostimulation of mOFC alcohol neuronal ensemble projections to the MD during T1 (Fig. 5c) significantly decreased the total amount of alcohol intake and the associated BEC. This result indicates that the role of mOFC→MD alcohol neuronal ensemble projections is sufficient to suppress alcohol consumption (Fig. 5d), without altering locomotor activity (Fig. 5e). In the real-time place preference assay, it induced a trend, indicating that the activation of this circuit may be aversive (Fig. 5f). Importantly, this effect was exclusive for alcohol intake since photoactivation of the mOFC→MD neuronal ensemble projections did not modify water, quinine or sacch consumption (Extended Data Fig. 11a-d). Finally, NpHR-mediated photoinhibition of the mOFC alcohol TRAPed neuronal terminals in the MD (Fig. 5g-i) produced a trend to increase alcohol consumption with no alteration in BEC (Fig. 5j), locomotor activity (Fig. 5k), time spent in a photoinhibited chamber in a real-time place preference assay (Fig 5l) neither water, quinine nor sacch consumption (Extended Data Fig. 11e-h). These results indicate that selective projections from the mOFC alcohol neuronal ensemble to the MD represent a key circuit to limit alcohol consumption.

**Fig. 5.**
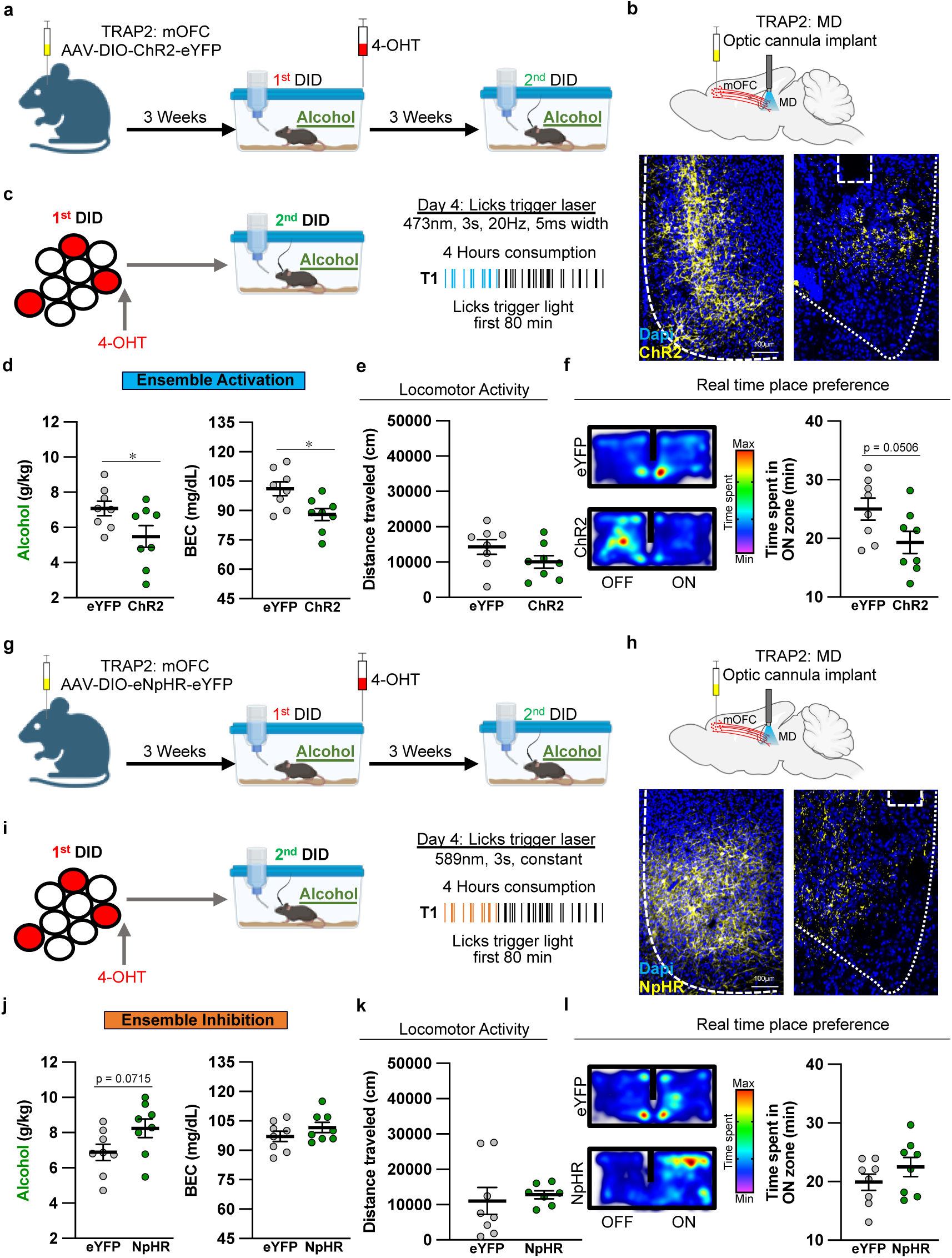
mOFC-MD photoactivation is sufficient to suppress binge alcohol drinking. **a.** Timeline of optogenetics experiments expressing ChR2 in TRAPed neurons in the mOFC and photostimulating their axons in the MD during binge alcohol drinking consumption. **b.** Representative images of the expression of ChR2-eYFP in the mOFC, axons from the mOFC and cannula trace in the MD. Broken white lines shows the boundaries of the mOFC, MD and the optic fiber trace. **c**. Pattern of light delivered to activate TRAPed neurons during the last day of the second DID. Blue light stimulation (473 nm, 20 Hz and 5 ms width) is triggered when mice establish contact with alcohol bottle spouts. Optogenetic stimulation has been delivered in T1. **d.** Alcohol consumption (*t*_(14)_= 2.14, *p*=0.04; *n*=8,8) and BEC measured the last day of the second DID during photostimulation (*t*_(14)_= 2.82, *p*=0.01; *n*=8,8). **e.** Locomotor activity of mice during photostimulation (*t*_(14)_= 1.56, *p*=0.14; *n*=8,8). **f.** Real-time preference/avoidance test of mice during photostimulation (*t*_(14)_= 2.13, *p*=0.05; *n*=8,8) and representative images. **g.** Timeline of optogenetics experiments expressing NpHR in TRAPed neurons in the mOFC and photoinhibiting the axons in the MD during binge alcohol drinking consumption. **h.** Representative images of the expression of NpHR-eYFP in the mOFC, axons from the mOFC and cannula trace in the PAG. Broken white lines shows the boundaries of the mOFC, MD and the optic fiber trace. **i**. Pattern of light delivered to inhibite TRAPed neurons during the last day of the second DID. Red light stimulation (589 nm, 3s, constant) is triggered when mice establish contact with alcohol bottle spouts. Optogenetic inhibition has been delivered in T1. **j.** Alcohol consumption (*t*_(14)_= 1.95, *p*=0.07; *n*=8,8) and BEC measured the last day of the second DID during photoinhibition (*t*_(14)_= 1.21, *p*=0.24; *n*=8,8). **k.** Locomotor activity of mice during photoinhibition (*t*_(13)_= 0.42, *p*=0.68; *n*=8,7). **o.** Real-time preference/avoidance test of mice during photoinhibition (*t*_(14)_= 1.2, *p*=0.24; *n*=8,8) and representative images. In all plots and statistical tests *n* represent biologically independent animals, *n* in caption represents biologically independent animals. Summary graphs show mean ± s.e.m.

## Discussion

The present study revealed a novel neuronal ensemble in the mOFC that is both necessary and sufficient to suppress alcohol consumption. We demonstrated that the formation of the mOFC neuronal ensemble depends on blood alcohol concentration reaching intoxication levels. Moreover, this neuronal ensemble is specific for alcohol consumption mirroring previous findings showing a neuronal ensemble in ventromedial prefrontal cortex that encoded cocaine seeking but not food seeking in rats^31^. Overall, these results suggest that different neuronal ensembles encode different stimuli instead of sharing information. Furthermore, we provided evidence that the stability of the alcohol neuronal ensemble across episodes of consumption lasted weeks, consistent with reports in other cortical areas^32^, indicating a persistent role of the neuronal ensemble in moderating binge drinking. Remarkably, our study is the first to report a neuronal ensemble that act as a sensor for intoxicating levels of alcohol in the brain after alcohol crosses the blood brain barrier^33^, the mechanisms underpinning this sensitivity remain undetermined. In parallel, we showed that sacch, a rewarding substance that does not produce addiction, induced the formation of a neuronal ensemble within the same region. Intriguingly, despite sharing similar molecular and cellular characteristics, alcohol and sacch neuronal ensembles are independent of each other and play different functional roles, suggesting that, while sharing similarities, neuronal ensembles may exhibit starkly different influence on behavior. Although the underlying cellular mechanisms driving neuronal ensemble formation remain elusive^34^ our findings point to AMPA receptors alterations and changes in calcium dynamics as a putative candidate involved in neuronal ensemble formation. Another surprising finding is the predominance of projecting GABAergic neurons in our neuronal ensemble, reminiscent of previous studies that identified similar long-range projecting GABAergic neurons in various regions of the neocortex^35,36^. Our study also revealed widespread projections throughout the brain, of the alcohol neuronal ensemble, unlike those of the sacch neuronal ensemble that were limited to a few regions. This aligns with recent findings on memory-related neuronal ensembles projecting widely^18^. The differential projectome between the alcohol and sacch neuronal ensembles may be key in determining the role of each neuronal ensemble. Although we identified the PAG, along with the MD, as the area receiving the densest innervation from the mOFC neuronal ensemble, we found that the mOFC→PAG ensemble circuit does not moderate binge alcohol drinking. This contrast recent findings showing that the medial prefrontal cortex to PAG pathway is associated with compulsive alcohol consumption^29^.

Although both results demonstrate the importance of the prefrontal cortex in controlling alcohol consumption, our data highlights the multifaceted roles that selective neuronal ensembles in this region could play in mediating various aspects of alcohol abuse. Importantly, in the present study we reveal that the mOFC neuronal ensemble circuit to the MD suppresses binge alcohol drinking. This is the first evidence of the role that this circuit plays in regulating alcohol consumption. The effect of the mOFC alcohol neuronal ensemble suppressing binge drinking represents the first instance of a neuronal ensemble acting as a suppressor of alcohol consumption, a novel insight in the addiction research landscape that offers promising avenues for the development of mOFC neuronal ensemble-targeted interventions.

## Methodology

### Animals

All experiments were performed following the guidelines for care and use of laboratory animals provided by the National Research Council. All the animal protocols involved in the manuscript were reviewed and approved by the Institutional Animal Care and Use Committee of the University of Massachusetts Chan Medical School (UMCMS). C57Bl/6J mice (Stock #000664, Jackson), TRAP2 mice (Stock #030323, Jackson), R26AI14/+ (AI14) mice (Stock #007914, Jackson) and GAD2Cre (Stock #10802, Jackson) were bred in the UMMS animal facility. Mice were genotyped through Transnetyx (Cordova, TN). TRAP2 mice were crossed with Ai14 mice to obtain the double heterozygous (TRAP2xAi14) mice that were used for the experimental work except when indicated otherwise. Approximately equal numbers of males and females were used in all the experiments except when we indicated that we are working with one or another sex. All mice were housed together and kept on a standard 12 h light/dark cycle (lights ON at 7 A.M.) with ad libitum access to food and water. Three to four weeks before experimentation, mice were kept under a reverse 12 h light/dark cycle (lights ON at 7 P.M.). All experiments were performed during the dark cycle phase (8 A.M. to 5 P.M.).

### Viral preparations

Biosensors, optogenetic and control plasmids packaged into viral particles (VP) were purchased from Addgene. For fiber photometry experiments we used pAAV.CAG.Flex.GCaMP6m.WPRE.SV40 (#100839-AAV5; titer ≥ 7×10¹² vg/mL) For optogenetic experiments, we used pAAV-EF1a-double floxed-hChR2(H134R)-EYFP-WPRE-HGHpA (#20298-AAV5; titer ≥ 1×10¹³ vg/mL), pAAV-Ef1a-DIO eNpHR 3.0-EYFP (#26966-AAV5; titer ≥ 1×10¹³ vg/mL) and pAAV-Ef1a-DIO EYFP (#27056-AAV5; titer ≥ 1×10¹³ vg/mL). For tracing experiments, we used pAAV hSyn FLEx mGFP-2A-Synaptophysin-mRuby (#71760-AAV1; titer ≥ 7×10¹² vg/mL). For experiments involving depletion of the neuronal ensemble, we used pAAV-flex-taCasp3-TEVp (#45580-AAV5; titer ≥ 7×10¹² vg/mL)

### Drug preparations

4-hydroxytamoxifen (4-OHT; Sigma, Cat# H6278) was dissolved at 20 mg mL^−1^ in ethanol by shaking at 37° C for 15 min and was then aliquoted and stored at –20° C for up to two months, as previously described^18^. Before use, 4-OHT was redissolved in ethanol by shaking at 37° C for 15 min, a 1:4 mixture of castor oil:sunflower seed oil (Sigma, Cat# 259853 and S5007) was added to give a final concentration of 10 mg mL^−1^ 4-OHT, and the ethanol was evaporated under vacuum by centrifugation. The final 10 mg mL^−1^ 4-OHT solutions were used immediately after preparation. All injections were delivered intraperitoneally (i.p.). Pyrazole (Sigma, Cat# P56607) was dissolved in saline and injected i.p. in a volume of 10 ml·kg^−1^ at a dose of 1 mmol·kg^−1^, as previously described ^37^. All injections were performed intraperitoneally (i.p.).

### Stereotaxic surgeries

Surgeries were performed under aseptic conditions as previously described^38^. Mice (4-5 weeks old) were deeply anaesthetized via intraperitoneal (ip) injection of a 100 mg/kg ketamine (VEDCO, Cat# 789-484-62) and 10 mg/kg xylazine (LLOYD, Cat# 789-161-65) mixture. Following anesthesia, the surgical area was shaved and disinfected with iodine and alcohol. Mice were then placed on a stereotaxic frame (Stoelting Co.) and ophthalmic ointment (Dechra, Cat# P1490-1) was applied to maintain eye lubrication. We used bregma and lambda landmarks to level the skull along the coronal and sagittal planes. A 0.4-mm drill was used for craniotomies at the target Bregma coordinates. For viral injections, mice were microinjected at a controlled rate of 50 nl/min using a gas-tight 33G 10-μl neurosyringe (1701RN; Hamilton) in a microsyringe pump (Stoelting Co). After injection, the needle remained unmoved for 5 min before a slow withdrawal. We performed injections in mOFC following the next coordinates: +2.25 AP, 0.33 ML and -2.5 DV. For tracing and photometry experiments mice were injected in one hemisphere and for casp3 and optogenetic experiments mice were injected bilaterally, with a viral volume of 300 nl. For fiber photometry or optogenetic experiments, we placed an optic fiber implant 2 weeks post viral injection. The optic fiber was placed above the injection site of the area of interest: mOFC (+2.25 AP, 0.33 ML and -2.4 DV; Doric Lenses, Cat# DFC_200/250-0.66_2.5mm_GS1.0_FLT for optogenetics and Cat# MFC_200/250-0.66_2.5mm_SMR_FLT for photometry); PAG (−3.08 AP, 3.00 ML and 0.25 DV; Doric Lenses, Cat# DFC_200/250-0.66_3mm_GS0.5_FLT) and MD (−1.58 AP, 3.00 ML and 0.33 DV; Doric Lenses, Cat# DFC_200/250-0.66_3mm_GS0.7_FLT). All optic fiber implants were secured into the skull using dental cement (C&B Metabond cement, Parkell Inc, Cat# S380). After each surgery, mice received IP injections of 1 mg/kg ketoprofen analgesic (Zoetis) and monitored post-surgery. Mice were allowed to recover in their home cages for 1 week before DID. Injection sites and viral expression were confirmed for all animals by experimenters blinded to behavioral outcome. Animals showing no viral or off target site viral expression or incorrect optic fiber placement (< 10%) were excluded from analysis.

### Drinking in the Dark Procedure

The paradigm “drinking in the dark” (DID) was used as a model of binge drinking^16,39^. Mice were individually housed and habituated for 7 days to drinking water ad libitum from a 25 mL serological pipette fitted with a drinking spout. For 4 consecutive days, starting 3 h after lights-off, water was replaced by alcohol 20% (v/v), for 2 h during the first 3 days and for 4 h on the 4^th^ day. Any alterations of this general procedure (i.e changing alcohol 20% for alcohol 10%) was indicated in the correspondent experiment. Body weights of mice were recorded weekly while alcohol, sacch or water intake values were recorded daily. These data were used to calculate the self-administered dose (i.e., g/kg). Control mice were always exposed to sacch (0.03%) or water. This protocol led to plasma alcohol levels of 117.5 ± 19.7 mg/dL immediately after 4 h of alcohol exposure (intoxication level as definided by NIAAA).

### Electrophysiological recordings

#### Slice preparation

We prepared coronal slices from fresh brain tissue of TRAP2xAi14 mice 10 days after the DID episode of consumption as previously described^40,41^. Following intracardiac perfusion with an ice-cold N-methyl-D-glucamine-based solution (in mM; 92 N-methyl-D-glucamine (NMDG), 2.5 KCl, 1.25 NaH_2_ PO_4_.H_2_O, 30 NaHCO_3_, 20 HEPES, 25 D-Glucose, 2 thiourea, 5 Na^+^ -ascorbate, 3 Na^+^ - pyruvate, 0.5 CaCl_2_, 10 MgSO_4_.7H_2_O) we rapidly removed and transferred the brain to a cold (∼ +0.5°C) oxygenated (95% O_2_ and 5% CO_2_) same N-methyl-D-glucamine-based cutting solution. Slices (200 μm) were cut with a Vibroslicer (VT1200, Leica MicroInstruments; Germany) and immediately transferred in an incubation chamber where they were left to recuperate in the NMDG-based solution for 20 min at 30 °C before being moved into a chamber containing artificial cerebrospinal fluid (aCSF; in mM): 120.6 NaCl, 0.25 KCl, 0.13 NaH_2_PO_4_ .H_2_O, 0.1 MgCl_2_ , 0.2 CaCl_2_, 26 NaHCO_3_ , 10 D-Glucose, at room temperature. Slices were left in this chamber for at least one hour before being placed in a recording chamber and perfused with ACSF at a constant rate of 2–3 ml/min at room temperature (∼21 °C).

#### Electrophysiology

We performed whole-cell patch clamp recordings as previously described^40,41^. When recording action potentials, we filled borosilicate glass electrodes (1.5 mm OD, 4-6 MΩ resistance) with an internal solution containing (mM): 120 K-methanesulfonate; 20 KCl; 10 Hepes; 2 K_2_ATP, 2 K_2_GTP, and 12 phosphocreatine. We replaced K-with Cs-Methanesulfonate and added 1 µM QX-314 (Ascent Scientific, Cambridge, MA USA) in the pipette internal solution when recording AMPA and NMDA EPSCs. Following seal rupture, series resistance (Rs), typically ranging between 10 and 20 MΩ, was fully compensated in current-clamp recording mode, and periodically monitored throughout recording sessions. Recordings with Rs changes larger than 20% were rejected. We acquired voltage and current traces in whole-cell patch-clamp with an EPC10 amplifier (HEKA Elektronik; Germany). We sampled and filtered voltage and current traces with PatchMaster 2.15 (HEKA Elektronik; Germany) at 10 kHz (20 kHz during induction of plasticity), and 2 kHz respectively. We subsequently analyzed all traces off-line using FitMaster 2.15 (HEKA Electronik; Germany). We evoked EPSPs by placing a concentric bipolar stimulating electrode in mOFC layer 2 and delivered brief (0.5 ms) electric pulses. When examining AMPA / NMDA ratio in voltage-clamp mode, we evoked AMPA-EPSCs at -70 mV in presence of 15 µM bicuculline (BIC) (Sigma-Aldrich; Saint Louis, MO USA). After blocking AMPA-EPSPs with 5 µM NBQX (Sigma-Aldrich; Saint Louis, MO USA) perfusion, we held membrane potential at +40 mV to evoke NMDA-EPSCs.

### Immunostaining and microscopy

Immunostaining and microscopy were performed as described previously^38^ with minor modifications. Mice were euthanized with sodium pentobarbital (200 mg/kg) before being transcardially perfused with ice-cold 0.1 M Sodium Phosphate buffer (pH 7.4). This was followed by 25 ml of cold 4% (W/V) paraformaldehyde (PFA) in 0.1 M Sodium Phosphate buffer. Brains were post-fixed for 24h in 4% PFA and transferred to a 20 % sucrose Sodium Phosphate buffer for 48h. Coronal sections (20 μm) were obtained using a freezing microtome (HM430; Thermo Fisher Scientific, MA, USA).

For immunofluorescence experiments brain sections were permeabilized with 3 washes of 0.1 M PBS for 10 min, blocked with 10 % donkey serum in 0.1 M of PBS with 0.5% Triton X-100 (Sigma, Cat# X100) and then incubated overnight with the corresponding primary antibody in 0.1 M of PBS with 0.5% Triton X-100 (Sigma, Cat# X100). Primary antibodies used: mouse anti-cFos 1:500 (Abcam, Cat# ab208942), chicken anti-GFP 1:500 (Abcam, Cat# ab13970), rabbit anti-GAD65 1:250 (Abcam, Cat# ab239372) and rabbit anti-tdTomato 1:250 (Rockland, Cat# 600-401-379). The neuronal ensemble was visualized using the endogenous fluorescence of the TRAP2xAi14 mice. Slices were subsequently washed for 3 times in 0.1 M PBS with 0.5% Triton X-100 during 5 min and incubated in secondary antibody in 0.1 M PBS with 0.5% Triton X-100 for 2 h. Secondary antibodies used: Alexa Fluor 488 goat anti mouse (Thermo Fisher, Cat# A11029); Alexa Fluor 488 goat anti chicken (Thermo Fisher, Cat# A11039), Alexa Fluor 488 goat anti rabbit (Thermo Fisher, Cat# A11008) and Alexa Fluor 488 donkey anti rabbit (Thermo Fisher, Cat# A21206). After 3 washes in 0.1 M PBS for 10 min, nuclei were counterstained with DAPI (Thermo Fisher, Cat# 62248), sections were mounted, air-dried and coverslipped using Fluoromount^TM^ Aqueous Mounting Medium (Sigma, Cat# F4680). All slices were imaged on a Zeiss LSM700 scanning confocal microscope equipped with 405nm, 488nm, 555nm, and 639nm lasers and ZEN acquisition software (Zeiss; Oberkochen, Germany)

### Blood alcohol content measurement

BEC has been measured as previously described^39^. Briefly, blood samples were collected in heparinized syringes from a small incision in the tail with the animal anesthetized. We collected between 10 and 20 μl for each mouse. The samples were centrifuged at 1300 × g for 10 min in heparinized microcentrifuge tubes (10% of the total volume) and plasma was injected into an alcohol analyzer (AM1, Analox, UK). The rationale of the method consists of alcohol being oxidized by the enzyme alcohol oxidase in the presence of molecular oxygen. Therefore, the rate of oxygen consumption is directly proportional to the alcohol concentration. Plasma alcohol levels were calculated as mg/dL, using ethanol 100 mg/dL as standard.

### iDISCO+ Imaging

#### iDISCO+ Sample processing

We performed iDISCO+ following the protocol created by the authors of the technique. This protocol is under constant modifications and updates can be found at: http://www.idisco.info. Briefly, we perfused the mice transcardially with sterile PBS followed by 4% PFA in sterile PBS. Samples were postfixed overnight in 4% PFA in sterile PBS and then processed with the iDISCO+ immunolabeling protocol^27^. Samples were stained using rabbit anti-tdTomato 1:250 (Rockland, Cat# 600-401-379) as a primary antibody and Alexa Fluor 488 donkey anti rabbit (Thermo Fisher, Cat# A21206).

#### iDISCO+ imaging

One week after clearing, iDISCO+ samples were imaged by a blind experimenter on a light-sheet microscope (Ultramicroscope II, LaVision Biotec) equipped with a sCMOS camera (Andor Neo) and a 2×/0.5 NA objective lens (MVPLAPO 2×) with a working distance of 6 mm. Images acquired beyond -4 mm from Bregma were discarded for the posterior image processing. We imaged using 488 nm filter. The samples were scanned with a step-size of 4 μm obtaining around 1500 TIFF images per sample.

#### iDISCO+ image processing and analysis

The analysis was performed using SmartAnalytics (LifeCanvas Technologies). Briefly, TIFF images were down-sampled and aligned to the Allen Brain Atlas in a registration in two steps: A first automatic step and a second manual registration to refine the automatic registration performed in the first step. After the integration with the Allen Brain Atlas, the software determined a fluorescence intensity signal for each brain region. Heat-maps were performed in LifeCanvas based on the values obtained.

### Fiber photometry

We performed *in vivo* fiber photometry as previously described^38^. Fluorescent signal from a genetically encoded calcium sensor (GCaMP6s) were recorded using a Doric Instruments Fiber Photometry System. We used a LED driver to deliver excitation light at 465 nm (∼8.5 mW output) and 405 nm (∼5 mW output, isosbestic wavelength for the calcium sensor). Light was reflected into a 200 μm 0.66 N.A. optic fiber patch cord via the Dual Fluorescence Minicube. Emissions were detected with a femtowatt photoreceiver (Model 2151, Newport) and were amplified by transimpedance amplification to measure voltage output. Sampling (12 kHz) and lock-in demodulation of the fluorescence signals were controlled by Doric Neuroscience Studio software with a decimation factor of 50. A Doric behavior camera was connected to the Doric Neuroscience Studio software using USB 3.0 Vision interface to synchronize the photometry recordings with time-locked videos of alcohol consumption. Sip start and end times were scored from the time-locked videos in a blinded fashion.

Demodulated fluorescence signals were processed using Python. Signals were smoothed using a 3 Hz Butterworth lowpass filter. The 405 nm channel was scaled to the 465 nm channel using least squares linear regression. Δ*F*/*F_0_* was calculated from the scaled signals using Δ*F/F*^0^ = (465 nm signal – fitted 405 nm signal) / fitted 405 nm signal. Z-scores for sipping events were calculated using Δ*F/F*^0^ values from 3 s prior to the sip initiation as baseline. Maximum (peak fluorescence) and mean z-scores were calculated for each baseline and event and then averaged for each mouse for statistical analysis.

### Single-cell RNA sequencing

#### Dissociation of mouse mOFC to a single cell suspension

We perfused the mice using the same aCSF buffer as described for Electrophysiology. The mOFC from sacch and alcohol mice was dissected out of the mouse brain and collected in Dulbecco’s phosphate buffered saline (DPBS) supplemented with 1g/l D-glucose, 36 mg/l sodium pyruvate (Gibco). We used a pool of 5 brains of each condition per sample. The tissue was briefly rinsed in DPBS and to generate a single cell suspension, dissociation was performed on a Singulator 100 (S2 Genomics) following the manufacturer’s recommended protocol for mouse brain dissociation using the mouse brain enzyme reagent (S2 Genomics) resuspended in DMEM (Gibco). The rinsed samples were loaded onto a cell isolation cartridge (S2 Genomics) and loaded onto the instrument with the preconfigured “Mouse Brain Cells” program. On completion of the run, the cartridge was retrieved from the instrument, punctured with a 5 mL serological pipette and the cell suspension collected from the output chamber. The sample was centrifuged at 300 g for 5 min at 4°C in a swinging bucket rotor. The cell suspension was resuspended in DPBS supplemented with 0.04% bovine serum albumin (BSA) (Sigma). Debris removal and clean up was performed as previously described ^42^ and after centrifugation, the cell pellet was resuspended in DPBS supplemented with 0.04% BSA and filtered through a 30 μm MACS SmartStrainer (Miltenyi Biotec) cell strainer. Cells were counted in a Countess II FL (Invitrogen) using trypan blue stain (Invitrogen) and on achieving a cell viability of at least 70%, the samples were prepared for Gel Beads-in-emulsion (GEMs) (10x Genomics) generation and barcoding.

#### Generation of 10x Genomic library for RNA-Seq

The protocol followed has been previously reported ^42^. Briefly, to generate GEMs, a single cell suspension and the Chromium Next GEM Single Cell 3’ v3.1 kit (10x Genomics) were used under the manufacturer’s recommended guidelines. To barcode the single cells, the gel beads together with the cell suspension in master mix and partitioning oil were loaded onto a Next GEM Chip G (10x Genomics) and ran on the Chromium Controller (10X Genomics). The resulting GEMs were transferred from the chip into pre-chilled tube strips and reverse transcribed to generate cDNA. For quality control and quantification, the cDNA was run on a Bioanalyzer High Sensitivity chip (Agilent) and Qubit dsDNA HS assay (Invitrogen) respectively. The cDNA was used to generate 3’ gene expression libraries using the Chromium Next GEM Single Cell 3’ Library kit v3.1 (10x Genomics) which were then shipped for sequencing (Novogene Inc) after post library construction quality control with a Bioanalyzer (Agilent).

#### Analysis

Cell Ranger (10x Genomics) was used to aggregate biological replicates, perform alignment, filter, count barcodes and UMIs. To identify cell types that underwent recombination as a result of Fos activity in TRAP2xAi14 mice, the sequence corresponding to the post recombination product placing one loxP site and the Frt promoter adjacent to one another was appended to the mm10 reference genome using Cell Ranger’s mkref function. This sequence is as follows: 5’-CATTTTGGCAAAGAATTGATTTGATACCGCGGGCCCTAAGAAGTTCCTATTCTCTAG AAAGTATAGGAACTTCGTCGACATTTAAATCATTTAAATATAACTTCGTATAATATA ACTTCGTATAATGTATGCTATACGAAGTTATTAGGTCCCTCGACCTGCAGCCCAAGC TAGT-3’ ^43^

Clustering, differential gene expression analysis and integrative analysis of control and stress conditions were performed using the Seurat V4 package ^44^. Low quality cells (genes < 200 or genes > 7500, or percentage of mitochondrial genes > 5%) were removed prior to analysis. 38,960 cells across alcohol and sacch samples were then used for downstream analysis. Briefly, gene counts were scaled by the cellular sequencing depth (total UMI) with a constant scale factor (10,000) and then natural-log transformed (log1p). 2,000 highly variable genes were selected in each sample based on a variance stabilizing transformation. Anchors between individual data were identified and correction vectors were calculated to generate an integrated expression matrix, which was used for subsequent clustering. Integrated expression matrices were scaled and centered followed by principal component analysis (PCA) for dimensional reduction. PC1 to PC20 were used to construct nearest neighbor graphs in the PCA space followed by Louvain clustering to identify clusters (resolution = 0.5). For visualization of clusters, Uniform Manifold Approximation and Projection (UMAP) was generated using the same PC1 to PC20. Cell clusters were manually annotated according to the expression of widely used cell-type specific gene markers ^45^. Expression of *Meg3*, *Thy1*, *Stmn2* and *Map2* identified neuronal clusters and these clusters were then analyzed for expression of the recombined sequence described above. As TRAPed neurons were predominantly expressed in Cluster Neuron_1 we subsequently used this cluster for further plots representations comparing TRAPed and Non TRAPed neurons.

### Behavior

#### Stress protocol

The experiment was performed during the dark cycle under dim red light as previously described^42^. We restrain mice for 5 min in a plastic restraint cone (Mouse Decapicone, Braintree Scientific INC; Cat# MDC-200). Two hours after the restraint, we injected 4-OHT to label the neuronal ensemble or we euthanized the mice to measure Fos.

#### Nicotine consumption

Mice had access to a two-bottle choice paradigm between water and nicotine as previously described ^46^. Every three days the concentration of nicotine was increased as follows (5, 10, 15, 20, 40, 60, 120, 240, 480 and 960 μg/ml). The last day of consumption depending on the group we injected 4-OHT to label the neuronal ensemble or we euthanized the mice to measure Fos.

#### Social Defeat

Mice in the defeated group were exposed to 4 episodes of SD during adulthood, each lasting 25 min and consisting of three phases as previously described^47^. We first introduced the “intruder” (the experimental animal) into the home cage of the “resident” (an aggressive opponent from the OF1 strain) for 10 min. During this initial phase, the intruder was protected from attack, but the wire mesh walls of the cage allowed for social interactions and species-typical threats from the male aggressive resident, thus facilitating instigation and provocation. In the second phase, the wire mesh was removed from the cage to allow confrontation between the two animals over a 5 min period. Finally, the wire mesh was put back in the cage to separate the two animals once again for a further 10 min to allow for social threats by the resident. Intruder mice were exposed to a different aggressor mouse during each SD episode. The criterion used to define an animal as defeated was the adoption of a specific posture signifying defeat, characterized by an upright submissive position, limp forepaws, upwardly angled head, and retracted ears. The last day of the protocol depending on the group we injected 4-OHT to label the neuronal ensemble or euthanized the mice to measure Fos.

#### Social exploration

Mice were tested in an open-field apparatus containing two plastic cylinders on opposite corners of the maze as previously described^48^. Experimental mice were subjected to a 5 min habituation session followed by a 5 min test session in which a juvenile mouse (C57BL/6J) was placed inside one of the two cylinders (counterbalanced). The experimental mouse was allowed to freely explore the field. 2 h after the encounter depending on the group, we injected 4-OHT to label the neuronal ensemble or sac the mice to measure Fos.

### Optogenetic experiments

#### Drinking in the Dark

For optogenetics experiments, we used the same DID protocol as described above. For photostimulation experiments, we used 10 mW of 473 nm light delivered in 5ms pulses at 20 Hz and for photoinhibition experiments, we used 5 mW of 589 nm light delivered continuously. In this experiment the serological pipette that delivered alcohol was linked to a custom-made lickometer based on Arduino. This lickometer was connected to the optogenetic equipment (Doric Lenses) and licks triggered the optic patter for 3 s after the lick. Mice were optogenetically stimulated the last day of consumption of the DID. To avoid damage to the neurons, we split the period of drinking consumption in three periods (T1, T2 and T3) and each mouse only received photostimulation/photoinhibition during one of those three periods.

#### Real-time place preference/avoidance

We used a real-time place preference/avoidance assay to determine if time spent in one chamber was associated with stimulation of the neuronal ensembles as previously described^29^. Mice were restrained and fiber optic patch cables were attached to the implanted ferrules before being placed in a plexiglass arena (24 in (l) × 10 in (w) × 20 in (h)) and allowed to move freely between two compartments for 45 minutes. Entry into the conditioned side of the chamber trigged a laser on period, which lasted as long as the animal remained on that side. For photostimulation experiments we used 10mW of 473nm light delivered in 5ms pulses at 20 Hz and for photoinhibition experiments we used 5mW of 589nm light delivered continuously. Mice were tested on two consecutive days, and on the second day the stimulation side and no stimulation side were reversed (order of which side was on/off first was counterbalanced across animals). A video camera positioned directly above the arena tracked and recorded movement using EthoVision XT 11.5 (Noldus Apparatus). All data presented are tracked from the ‘center’ of the subject, and time spent in each zone was averaged across the two testing sessions.

#### Open Field

We performed open field to measure locomotor activity. Mice were individually placed facing one of the walls of a Plexiglass open field (24 in (l) × 10 in (w) × 20 in (h)). Mice were allowed to explore freely for 30 min and the locomotor activity of the mice was automatically tracked using EthoVision XT 11.5 (Noldus Apparatus). The animals received light stimulation for the entire 30 min session with the same parameters as described for the real-time place preference/avoidance test.

### Statistical analysis and Reproducibility

Data were analyzed by means of two-tailed unpaired t-test, one-way or two-way ANOVAs with/without repeated-measures, as indicated. Tukey test was used as a post hoc for multiple comparisons. Grubbs’ test was applied to check for outliers. Comparisons of z-scores or AUC photometry signals were made using the calculated average for each animal. Each data set was tested for normal distribution prior to analysis and presented as mean ± standard error of the mean (SEM). All behavioral experiments were conducted in at least two independent cohorts of animals. Imaging analyses included representative pictographs of at least three animals included in the same experiment. All statistical analyses were performed in GraphPad Prism 10.1.1. Software (Graphpad Software Inc.) and statistical significance was established at p<0.05.

### Data availability

All data needed to evaluate the conclusions in the paper are present in the paper and/or the Supplemental Material. Data will be made available on request to the corresponding author. Custom codes used in this manuscript are publicly accessible on Harvard Dataverse using the following link: (https://doi.org/10.7910/DVN/XWKZ8N).

Single cell sequencing data are available on the NCBI submission portal (https://nam10.safelinks.protection.outlook.com/?%3A%2F%2Fdataview.ncbi.nlm.nih.gov%2Fobject%2FPRJNA1092677%3Freviewer%3Do21piokjenq4kst5kubfhjf1en&data=05%7C02%7CMax.Zinter%40umassmed.edu%7C39b75ead47bd4fc2005508dc52727214%7Cee9155fe2da34378a6c44405faf57b2e%7C0%7C0%7C638475898707765357%7CUnknown%7CTWFpbGZsb3d8eyJWIjoiMC4wLjAwMDAiLCJQIjoiV2luMzIiLCJBTiI6Ik1haWwiLCJXVCI6Mn0%3D%7C0%7C%7C%7C&sdata=XcUakhotDk7sYZ1nr%2F8a%2BPR8DOSClXd1NiZPzBA3SQY%3D&reserved=0).

## Acknowledgments.

We thank C. LaFlash for assistance in maintaining mouse lines, C. Baer for her support with whole-brain analysis and J Sternberg from Life Science Editors for editing support and BioRender. This work was supported by AA027807 (G.E.M), R01NS112492 (T.T), T32GM135751 (T.L), BrightFocus Foundation A2022006F and Alzheimer’s Association AARF-22-923219 (V.D.L), DA041482 and DA047678 (A.R.T), 2021 BBRF Young Investigator Grant-30616 (S.M), NIMH-R01MH113743 (D.P.S), NINDS-R01NS117533 (D.P.S) and NIA-RF1AG068281 (D.P.S).

## Author contributions

P.G., S.M., A.R.T., D.P.S., T.T., and G.E.M. contributed to the study design. P.G., T.L., M.Z., M.P., V.D.L., T.G.F., and G.E.M performed the experiments and analyzed the data. P.G., and G.E.M. prepared the manuscript. All authors contributed to editing and reviewing the manuscript.

## Competing Interest

All authors report no competing financial interests or potential conflicts of interest. The content is solely the responsibility of the authors and does not necessarily represent the official views of the National Institutes of Health.

**Extended Data Fig.1. Binge alcohol drinking elicits a strong Fos signal in the mOFC. a.** Schematic of the DID protocol. Following the same drinking pattern, mice have access to alcohol, water or sacch. **b.** Water consumption in males and females (*F*_day(3,36)_ = 20.32, *p* < 0.0001; *F*_sex(1,36)_ = 0.53, *p* = 0.47; *F*_interaction(3,36)_ = 0.22, *p* = 0.87; * *vs* Day 1 same group; *n*=5,6). **c.** Sacch consumption in males and females (*F*_day(3,36)_ = 26.87, *p* < 0.0001; *F*_sex(1,36)_ = 0.11, *p* = 0.73; *F*_interaction(1,36)_ = 0.67, *p* = 0.57; * *vs* Day 1 same group; *n*=5,6). **d.** Alcohol consumption (*F*_day(3,37)_ = 11.72, *p* < 0.0001; *F*_sex(1,37)_ = 0.79, *p* = 0.37; *F*_interaction(1,37)_ = 0.56, *p* = 0.64; * *vs* Day 1 same group. *n*=6,6) and BEC (*t*_(10)_= 0.37, *p*=0.71; *n*=6,6) in males and females. **d.** Timeline of brain perfusion to measure Fos expression the last day of DID. We euthanized mice at the end of the 4 h of consumption (0 h), and 1 h, 2 h and 6 h post drinking. **f.** Representative images of Fos^+^ neurons expression after water, sacch and alcohol consumption at the time points selected. The broken white lines show the medial boundary of the mOFC. **g.** Fos^+^ neurons (expressed as a % of the total DAPI cells) in water, sacch and alcohol drinking mice (*F*_group(2,60)_ = 17.81, *p* < 0.0001; *F*_time(3,60)_ = 32.4, *p* < 0.0001; *F*_interaction(6,60)_ = 5.66, *p* = 0.0001; * *vs* 0h; # *vs* Sacch 2h: *n*=6,6). **h.** Fos^+^ neurons (expressed as a % of the total DAPI cells) for water, sacch and alcohol groups 2 h post drinking in males and females (*F*_group(2,28)_ = 56.87, *p* < 0.0001; *F*_sex(1,28)_ = 0.34, *p* = 0.56; *F*_interaction(2,28)_ = 0.84, *p* = 0.44; * *vs* Water same sex; # *vs* Sacch same sex; *n*=5,5,6,6 5,6). **i.** Fos^+^ neurons per FOV in water, Sacch, and alcohol groups. In all plots and statistical tests, *n* represents biologically independent animals, *n* in caption represents biologically independent animals. Summary graphs show mean ± s.e.m.

**Extended Data Fig. 2. The alcohol neuronal ensemble is not involved in other risk behaviors associated with alcohol consumption. a.** Strategy to study the effect of binge alcohol drinking on the formation of neuronal ensembles activated by other behaviors (Stress, Nicotine consumption, social defeat and social exploration). **b.** Representative images of TRAPed neurons expression for the groups used. The broken white lines show the medial boundary of the mOFC. **e.** Strategy to study the effect of other behaviors in binge alcohol drinking and potential effects of the activity of other behaviors (Stress, Nicotine consumption, social defeat and social exploration) on the size of the alcohol TRAPed neurons. **c.** Quantification of percentage of TRAPed neurons that are Fos^+^ neurons (*F*_(4,30)_ = 80.28, *p* < 0.0001; *n*=7,7,7,7,7). **d.** Quantification of percentage of Fos^+^ neurons that are TRAPed neurons (*F*_(4,30)_ = 80.28, *p* < 0.0001; *n*=7,7,7,7,7). **e.** Strategy to study the effect of other behaviors in binge alcohol drinking and potential effects of the activity of other behaviors (Stress, Nicotine consumption, social defeat and social exploration) on the size of the alcohol TRAPed neurons. **f.** Representative images of TRAPed neurons expression for the groups used. The broken white lines show the medial boundary of the mOFC. **g.** TRAPed neurons (expressed as a % of the total DAPI cells) for the groups used (*F*_(5,30)_ = 0.8, *p* = 0.55; *n*=6,6,6,6,6,6). **h.** Alcohol consumption at the end of the last day of DID (*F*_(5,31)_ = 0.61, *p* = 0.69; n= 6,6,6,7,6,6). In all plots and statistical tests, *n* represents biologically independent animals, *n* in caption represents biologically independent animals. Summary graphs show mean ± s.e.m.

**Extended Data Fig. 3. The alcohol neuronal ensemble is exclusively activated during alcohol consumption. a.** Timeline of fiber photometry experiments in TRAPed neurons after binge alcohol drinking consumption while they drink water, sacch or quinine in a 2^nd^ DID episode (T3). **b.** Mean mOFC GCaMP-recorded calcium activity in TRAPed neurons 2 seconds prior to the sip (vertical broken line) and 4 seconds after the sip in a 2^nd^ DID in which the mice drinks water, sacch or quinine. Insert show heatmap raster plot of mOFC GCaMP6s-recorded calcium activity in TRAPed neurons in a representative mouse that drinks water, sacch or quinine. **c.** Peak Fluorescence during base (−2 to 0 seconds) and event (0 to 4 seconds) in mice that drink water (*t*_(64)_= 0.64, *p*=0.52; *n*=8). **d.** Peak Fluorescence during base (−2 to 0 seconds) and event (0 to 4 seconds) in mice that drink sacch (*t*_(56)_= 0.27, *p*=0.78; *n*=9). **e.** Peak Fluorescence during base (−2 to 0 seconds) and event (0 to 4 seconds) in mice that drink quinine (*t*_(50)_= 0.5, *p*=0.61; *n*=8). In all plots and statistical tests, *n* represents biologically independent animals, *n* in caption represents biologically independent animals. Summary graphs show mean ± s.e.m.

**Extended Data Fig. 4. Intoxicating levels of alcohol activate the alcohol neuronal ensemble. a.** Schematic of DID protocols in mice drinking 10 or 20% alcohol. **b.** Alcohol consumption (*F*_group(1,56)_ = 200.7, *p* < 0.0001; *F*_day(3,56)_ = 100.4, *p* < 0.0001; *F*_interaction(3,56)_ = 5.87, *p* = 0.0015; * *vs* 10% Day 4. *n*=8,8). **c.** BEC measured the last day of consumption (*t*_(14)_= 12.1, *p*<0.0001; *n*=8,8). **d.** Representative images of TRAPed neurons in mice drinking a 10 or 20% alcohol solution. The broken white lines show the medial boundary of the mOFC. **e.** TRAPed neurons (expressed as a % of the total DAPI cells) measured after alcohol consumption in mice drinking a 10 or 20% alcohol solution (*t*_(14)_= 11.91, *p*<0.0001; *n*=8,8). **f.** Schematic of the DID protocols where mice consumed a 20 % alcohol solution for either 4 h or 2 h on the last binge drinking day. In the latter, we injected pyrazole to the solution to prevent alcohol from being metabolized and increase BEC. **g.** Alcohol consumption (*F*_group(2,80)_ = 17.56, *p* < 0.0001; *F*_day(3,80)_ = 10.18, *p* < 0.0001; *F*_interaction(6,80)_ = 8.86, *p* < 0.0001; * *vs* 2h day 4; # *vs* 2h + pyr day 4. *n*=8,8,8). **h**. BEC measured the last day of DID (*F*_(2,21)_ = 29.30, *p* < 0.0001; * *vs* 4h; # *vs* 2h; n= 8,8,8). **i.** Representative images of TRAPed neurons expression for the groups used. The broken white lines show the medial boundary of the mOFC. **j.** TRAPed neurons (expressed as a % of the total DAPI cells) measured after alcohol consumption (*F*_(2,21)_ = 60.99, *p* < 0.0001; * *vs* 4h; # *vs* 2h; n= 8,8,8). **k.** Schematic of the DID protocol varying the duration of the last day of DID in mice that drinks sacch. **l.** Sacch consumption (*F*_group(1,52)_ = 42.12, *p* < 0.0001; *F*_day(3,52)_ = 67.78, *p* < 0.0001; *F*_interaction(3,52)_ = 34.99, *p* < 0.0001; * *vs* 2h day 4; *n*=7,8). **m.** Representative images of TRAPed neurons expression for the groups used. The broken white lines show the medial boundary of the mOFC. **n.** TRAPed neurons (expressed as a % of the total DAPI cells) measured after alcohol consumption (*t*_(13)_= 0.77, *p*=0.44; *n*=8,8) In all plots and statistical tests, *n* represents biologically independent animals, *n* in caption represents biologically independent animals. Summary graphs show mean ± s.e.m.

**Extended Data Fig. 5. The alcohol neuronal ensemble and non-TRAPed neurons show different active membrane properties. a.** Schematic of electrophysiological experiments. **b.** Current-voltage relationship of NMDA current between -60 and +40 mV. Values are expressed as percent of current evoked at -60 mV. EPSCs were evoked every 10 sec 8 consecutive times (*F*_group(1,102)_ = 5.74, *p* = 0.018; *F*_Mv(5,102)_ = 130.8, *p* < 0.0001; *F*_interaction(5,102)_ =0.79, *p* = 0.55; *n*=10,9). **c.** AMPA/NMDA current ratio (*t*_(18)_= 0.94, *p*=0.35; *n*=10,10) and representative traces. **d.** Dot plot illustrating scaled expression levels (color) and the proportions of expressing neurons (dot size) of genes encoding sodium and potassium channels. **e.** Representative I-V traces from non TRAPed neurons (black traces) and TRAPed neurons (green traces). **f.** RMP (*t*_(32)_= 0.11, *p*=0.9; *n*=9,9). **g.** AP amplitude (*t*_(32)_= 5.68, *p*<0.0001; *n*=9,9). **h.** Firing (*t*_(32)_= 5.08, *p*<0.0001; *n*=9,9). **i.** SAG (*t*_(32)_= 4.29, *p*=0.0002; *n*=9,9). **j.** Threshold (*t*_(32)_= 1.51, *p*=0.13; *n*=9,9). **k.** fAHP (*t*_(32)_= 1.49, *p*=0.14; *n*=9,9). **l.** Rin (*t*_(32)_= 0.37, *p*=0.71; *n*=9,9). **m.** AP width (*t*_(32)_= 1.88, *p*=0.068; *n*=9,9). **n.** Rheobase (*t*_(34)_= 0.02, *p*=0.98; *n*=9,9). In all plots and statistical tests *n* represent cells (except for **b** that represent the average of cells), *n* in caption represents biologically independent animals. Summary graphs show mean ± s.e.m.

**Extended Data Fig. 6. Sacch and alcohol neuronal ensembles show similar transcriptomical pattern and electrophysiological. a.** Feature plot showing normalized expression values for GAD2 across the TRAPed neurons in the UMAP space in sacch drinking mice. **b.** Representative images of sacch TRAPed neurons, GAD2^+^ neurons, overlap TRAPed/GAD2^+^ neurons and proportion of GAD2^+^ in the alcohol neuronal ensemble. **c.** Dot plot illustrating scaled expression levels (color) and the proportions of expressing neurons (dot size) of glutamate and GABA-receptor-related genes. **d.** Schematic of electrophysiology experiments. **e.** Representative images of mOFC neurons in DIC and 550nm conditions. **f.** Representative AMPA-EPSCs from sacch non TRAPed and TRAPed neurons recorded in the presence of NMDA and GABA_A_ receptor antagonists at holding potentials of -60 and +60mV. EPSCs were evoked every 10 sec 8 consecutive times. Average traces are shown in gray and orange for non TRAPed and TRAPed neurons, respectively. **g.** Current-voltage relationship of AMPA current between -60 and +60 mV. Values are expressed as percent of current evoked at -60 mV (*F*_group(1,89)_ = 28.51, *p* < 0.0001; *F*_Vh(6,89)_ = 616.7, *p* < 0.0001; *F*_interaction(6,89)_ =9.97, *p* < 0.0001; * *vs* Non TRAPed same time; *n*=8,6). **h.** Current-voltage relationship of NMDA current between -60 and +40 mV. Values are expressed as percent of current evoked at -60 mV. EPSCs were evoked every 10 sec 8 consecutive times (*F*_group(1,102)_ = 10.08, *p* < 0.0001; *F*_Mv(5,102)_ = 105.6, *p* < 0.0001; *F*_interaction(5,102)_ =0.59, *p* = 0.7; *n*=10,8). **i.** AMPA/NMDA current ratio (*t*_(18)_= 0.94, *p*=0.35; *n*=10,10) and representative traces. Orange traces represent averages of 8 consecutives EPSCs evoked every 10 sec. **j.** Dot plot illustrating scaled expression levels (color) and the proportions of expressing neurons (dot size) of genes encoding sodium and potassium channels. **k.** Representative I-V traces from non-TRAPed neurons (black traces) and TRAPed neurons (green traces). **l.** RMP (*t*_(43)_= 0.04, *p*=0.96; *n*=9,9). **m.** AP amplitude (*t*_(42)_= 3.3, *p*=0.002; *n*=9,9). **n.** Firing (*t*_(40)_= 3.12, *p*=0.0033; *n*=9,9). **o.** SAG (*t*_(41)_= 3.32, *p*=0.0019; *n*=9,9). **p.** Threshold (*t*_(41)_= 2.68, *p*=0.01; *n*=9,9). **q.** fAHP (*t*_(42)_= 0.32, *p*=0.74; *n*=9,9). **r.** Rin (*t*_(41)_= 1.24, *p*=0.22; *n*=9,9). **s.** AP width (*t*_(42)_= 5.63, *p*<0.0001; *n*=9,9). **t.** Rheobase (*t*_(30)_= 0.44, *p*=0.66; *n*=9,9). In all plots and statistical tests *n* represent cells (except for **g** and **h** that represent the average of cells), *n* in caption represents biologically independent animals. Summary graphs show mean ± s.e.m.

**Extended Data Fig. 7. Neither activation nor inhibition of alcohol and sacch neuronal ensembles alter the consumption of other substances. a.** Timeline of optogenetics experiments using ChR2 in TRAPed neurons in binge alcohol drinking consumption. During a second DID episode mice consumed a substance other than alcohol (i.e., water, quinine, saccharine). **b.** Preference for alcohol during photostimulation alcohol TRAPed neurons (*t*_(14)_= 4.15, *p*<0.001; *n*=8,8). **c.** Water consumption during photostimulation of alcohol TRAPed neurons (*t*_(14)_= 0.57, *p*=0.57; *n*=8,8). **d.** Quinine consumption during photostimulation of alcohol TRAPed neurons (*t*_(14)_= 0.21, *p*=0.83; *n*=8,8). **e.** Sacch consumption during photostimulation of alcohol TRAPed neurons (*t*_(14)_= 0.63, *p*=0.53; *n*=8,8). **f.** Timeline of optogenetics experiments using ChR2 in TRAPed neurons in binge saccharine drinking mice. During a second DID episode, mice consumed a substance other than sacch (i.e., water, quinine, alcohol). **g.** Preference for sacch during photostimulation of sacch TRAPed neurons (*t*_(14)_= 0.99, *p*=0.33; *n*=8,8). **h.** Water consumption during photostimulation of sacch TRAPed neurons (*t*_(14)_= 0.24, *p*=0.81; *n*=8,8). **i.** Quinine consumption during photostimulation of sacch TRAPed neurons (*t*_(14)_= 0.69, *p*=0.49; *n*=8,8). **j.** Alcohol consumption during photostimulation of sacch TRAPed neurons (*t*_(14)_= 0.52, *p*=0.6; *n*=8,7). **k.** Timeline of optogenetics experiments using NpHR in TRAPed neurons in binge alcohol drinking consumption. During a second DID episode, mice consumed a substance other than alcohol. **l.** Preference for alcohol during photoinhibition of alcohol TRAPed neurons (*t*_(14)_= 2.95, *p*=0.014; *n*=8,8). **m.** Water consumption during photoinhibition of alcohol TRAPed neurons (*t*_(14)_= 0.27, *p*=0.78; *n*=8,8). **n.** Quinine consumption during photoinhibition of alcohol TRAPed neurons (*t*_(14)_= 0.51, *p*=0.61; *n*=8,8). **o.** Sacch consumption during photoinhibition of alcohol TRAPed neurons (*t*_(14)_= 0.8, *p*=0.43; *n*=8,8). **p.** Timeline of optogenetics experiments using NpHR in TRAPed neurons after sacch consumption. During a second DID episode mice consumed a substance other than sacch. **q.** Preference for sacch during photoinhibition of sacch TRAPed neurons (*t*_(14)_= 0.21, *p*=0.83; *n*=8,8). **r.** Water consumption during photoinhibition of sacch TRAPed neurons (*t*_(14)_= 0.28, *p*=0.77; *n*=8,8). **s.** Quinine consumption during photoinhibition of sacch TRAPed neurons (*t*_(14)_= 0.07, *p*=0.94; *n*=8,8). **t.** Alcohol consumption during photoinhibition of sacch TRAPed neurons (*t*_(14)_= 1.5, *p*=0.15; *n*=8,8). **u.** Locomotor activity of mice during photostimulation of the sacch TRAPed neurons (*t*_(14)_= 1.07, *p*=0.29; *n*=8,8). **v.** Real-time preference/avoidance test of mice during photostimulation of sacch TRAPed neurons (*t*_(14)_= 0.11, *p*=0.91; *n*=8,8) and representative images. **w.** Locomotor activity of mice during photoinhibition of sacch TRAPed neurons (*t*_(14)_= 0.53, *p*=0.6; *n*=8,8). **x.** Real-time preference/avoidance test of during photoinhibition of sacch TRAPed neurons (*t*_(14)_= 0.27, *p*=0.79; *n*=8,8) and representative images. In all plots and statistical tests *n* represent biologically independent animals, *n* in caption represents biologically independent animals. Summary graphs show mean ± s.e.m.

**Extended Data Fig. 8. Depletion of the alcohol neuronal ensemble increases long term consumption. a.** Timeline of chemogenetics experiments using Casp3 in TRAPed neurons in alcohol or sacch drinking mice. **b.** Representative image of the site of injection. The broken white lines show the medial boundary of the mOFC and Hamilton syringe. **c**. Strategy to ablate TRAPed neurons in the second episode of DID. 4-OHT injections triggers the expression of Casp3 restricted to TRAPed neurons, leading to cell death. **d.** Fos^+^ neurons (expressed as a % of the total DAPI cells) during a second DID in mice with intact (No 4-OHT) or ablated TRAPed neurons (4-OHT) (*t*_(14)_= 7.26, *p*<0.0001; *n*=8,8) and representative images. **e.** Fos^+^ neurons (expressed as a % of the total DAPI cells) during a second episode of sacch consumption in mice with intact (No 4-OHT) or ablated TRAPed neurons (4-OHT) (*t*_(14)_= 7.59, *p*<0.0001; *n*=8,8) and representative images. **f.** Alcohol consumption during 1^st^ and 2^nd^ episode of DID after ablation of TRAPed neurons (No 4-OHT: *t*_(6)_= 0.008, *p*=0.99; *n*=8,8; 4-OHT: *t*_(6)_= 5.68, *p*=0.001; *n*=7,7). **g.** Sacch consumption during 1^st^ and 2^nd^ episode of DID after ablation of TRAPed neurons (No 4-OHT: *t*_(7)_= 1.25, *p*=0.25; *n*=8,8; 4-OHT: *t*_(8)_= 1.99, *p*=0.08; *n*=8,8). **h.** Water consumption after ablation of binge alcohol drinking TRAPed neurons (*t*_(14)_= 0.67, *p*=0.51; *n*=8,8). **i**. Locomotor activity after ablation of alcohol TRAPed neurons (*t*_(12)_= 1.41, *p*=0.18; *n*=7,7). **j.** Preference for alcohol after ablation of alcohol TRAPed neurons (*t*_(14)_= 2.38, *p*=0.03; *n*=8,8). k. Compulsive alcohol drinking after ablation of alcohol TRAPed neurons, shown by resistance to quinine adulteration of the alcohol solution (*F*_Quinine(4,70)_ = 6.14, *p* = 0.0003; *F*_ablation(1,70)_ = 79.78, *p* < 0.0001; *F*_interaction(4,70)_ =1.14, *p* = 0.34; * *vs* No 4-OHT; *n*=8,8). **l.** Long term consumption of alcohol after ablation of the neuronal ensemble (*F*_ablation(3,35)_ = 10.36, *p* < 0.0001; *F*_time(1,7)_ = 21.38, *p* = 0.0024; *F*_interaction(5,35)_ =3.6, *p* =0.0098; * *vs* Week 1, # *vs* same time No 4-OHT; *n*=8,8). In all plots and statistical tests, *n* represents biologically independent animals (except for **k** and **l** that represent the average of animals). Summary graphs show mean ± s.e.m.

**Extended Data Fig. 9. GAD2 neurons does not control binge alcohol drinking. a.** Timeline of optogenetics experiments using ChR2 in GAD2 mice after binge alcohol drinking consumption and pattern of light delivered to activate the GAD2 neurons during the last day of DID. Blue light stimulation (473 nm, 20 Hz and 5 ms width) is triggered when mice establish contact with alcohol bottle spouts. **b.** Representative image of the expression of ChR2-eYFP in the mOFC. The broken white lines show the medial boundary of the mOFC and the optic fiber trace. **c.** Alcohol consumption (*t*_(14)_= 1.56, *p*=0.14; *n*=8,8) measured the last day of DID while the mice receive photoinhibition. **d.** Locomotor activity of mice during photostimulation (*t*_(13)_= 0.62, *p*=0.54; *n*=8,7). **e.** Real-time preference/avoidance test of mice during photostimulation (*t*_(14)_= 0.72, *p*=0.47; *n*=8,8) and representative images. **f.** Timeline of optogenetics experiments using NpHR in GAD2 mice after binge alcohol drinking consumption and pattern of light delivered to inibit the GAD2 neurons during the last day of DID. Red light stimulation (589 nm, 3s, constant) is triggered when mice establish contact with alcohol bottle spouts. **g.** Representative image of the expression of NpHR-eYFP in the mOFC. The broken white lines show the medial boundary of the mOFC and the optic fiber trace.**h.** Alcohol consumption (*t*_(11)_= 0.19, *p*=0.84; *n*=7,6) measured the last day of DID while the mice receive photoinhibition. **i.** Locomotor activity of mice during photoinhibition (*t*_(14)_= 1.45, *p*=0.16; *n*=8,8). **j.** Real-time preference/avoidance test of mice during photoinhibition (*t*_(14)_= 0.87, *p*=0.39; *n*=8,8) and representative images.). In all plots and statistical tests, *n* represents biologically independent animals. Summary graphs show mean ± s.e.m.

**Extended Data Fig. 10. mOFC-PAG circuit do not controls binge alcohol drinking. a.** Timeline of optogenetics experiments expressing ChR2 in TRAPed neurons in the mOFC and photostimulating the axons in the PAG during binge alcohol drinking consumption. **b.** Representative images of the expression of ChR2-eYFP in the mOFC and cannula trace in the PAG. Broken white lines shows the boundaries of the mOFC, PAG and the optic fiber trace. **c**. Pattern of light delivered to activate TRAPed neurons during the last day of the second DID. Blue light stimulation (473 nm, 20 Hz and 5 ms width) is triggered when mice establish contact with alcohol bottle spouts. Optogenetic stimulation has been delivered in T1. **d.** Alcohol consumption (*t*_(14)_= 0.23, *p*=0.81; *n*=8,8) and BEC measured the last day of the second DID during photostimulation (*t*_(14)_= 0.31, *p*=0.75; *n*=8,8). **e.** Locomotor activity of mice during photostimulation (*t*_(14)_= 0.32, *p*=0.74; *n*=8,8). **f.** Real-time preference/avoidance test of mice during photostimulation (*t*_(14)_= 0.91, *p*=0.37; *n*=8,8) and representative images. **g.** Timeline of optogenetics experiments expressing NpHR in TRAPed neurons in the mOFC and photoinhibiting the axons in the PAG during binge alcohol consumption. **h.** Representative images of the expression of NpHR-eYFP in the mOFC and cannula trace in the PAG. Broken white lines shows the boundaries of the mOFC, PAG and the optic fiber trace. **i**. Pattern of light delivered to inhibit TRAPed neurons during the last day of the second DID. Red light stimulation (589 nm, 3s, constant) is triggered when mice establish contact with alcohol bottle spouts. Optogenetic inhibition has been delivered in T1. **j.** Alcohol consumption (*t*_(14)_= 0.07, *p*=0.94; *n*=8,8) and BEC measured the last day of the second DID during photoinhibition (*t*_(14)_= 0.07, *p*=0.94; *n*=8,8). **k.** Locomotor activity of mice during photoinhibition (*t*_(12)_= 1.77, *p*=0.1; *n*=6,8). **o.** Real-time preference/avoidance test of mice during photoinhibition (*t*_(14)_= 0.27, *p*=0.79; *n*=8,8) and representative images. In all plots and statistical tests *n* represent biologically independent animals, *n* in caption represents biologically independent animals. Summary graphs show mean ± s.e.m.

**Extended Data Fig. 11. Neither activation nor inhibition of the alcohol neuronal ensemble projections alter the consumption of other substances.** a. **Timeline of optogenetics** experiments expressing ChR2 in TRAPed neurons in the mOFC and photostimulating the projections in MD. During a second DID episode mice consumed a substance other than alcohol (i.e., water, quinine, saccharine). **b.** Water consumption during photostimulation of alcohol TRAPed neurons (*t*_(14)_= 0.62, *p*=0.54; *n*=8,8). **c.** Quinine consumption during photostimulation of alcohol TRAPed neurons (*t*_(14)_= 0.61, *p*=0.54; *n*=8,8). **d.** Sacch consumption during photostimulation of alcohol TRAPed neurons (*t*_(14)_= 0.08, *p*=0.93; *n*=8,8). **e.** Timeline of optogenetics experiments using ChR2 in TRAPed neurons in the mOFC and photostimulating the projections in MD. During a second DID mice consumed a substance other than alcohol (i.e., water, quinine, saccharine).. **f.** Water consumption during photoinhibition of sacch TRAPed neurons (*t*_(14)_= 0.26, *p*=0.79; *n*=8,8). **g.** Quinine consumption during photoinhibition of sacch TRAPed neurons (*t*_(14)_= 0.82, *p*=0.42; *n*=8,8). **h.** Sacch consumption during photoinhibition of sacch TRAPed neurons (*t*_(14)_= 0.1, *p*=0.91; *n*=8,8). In all plots and statistical tests *n* represent biologically independent animals, *n* in caption represents biologically independent animals. Summary graphs show mean ± s.e.m.

