## Supplementary Figures for "An orbitocortical-thalamic circuit suppresses binge alcohol-drinking"

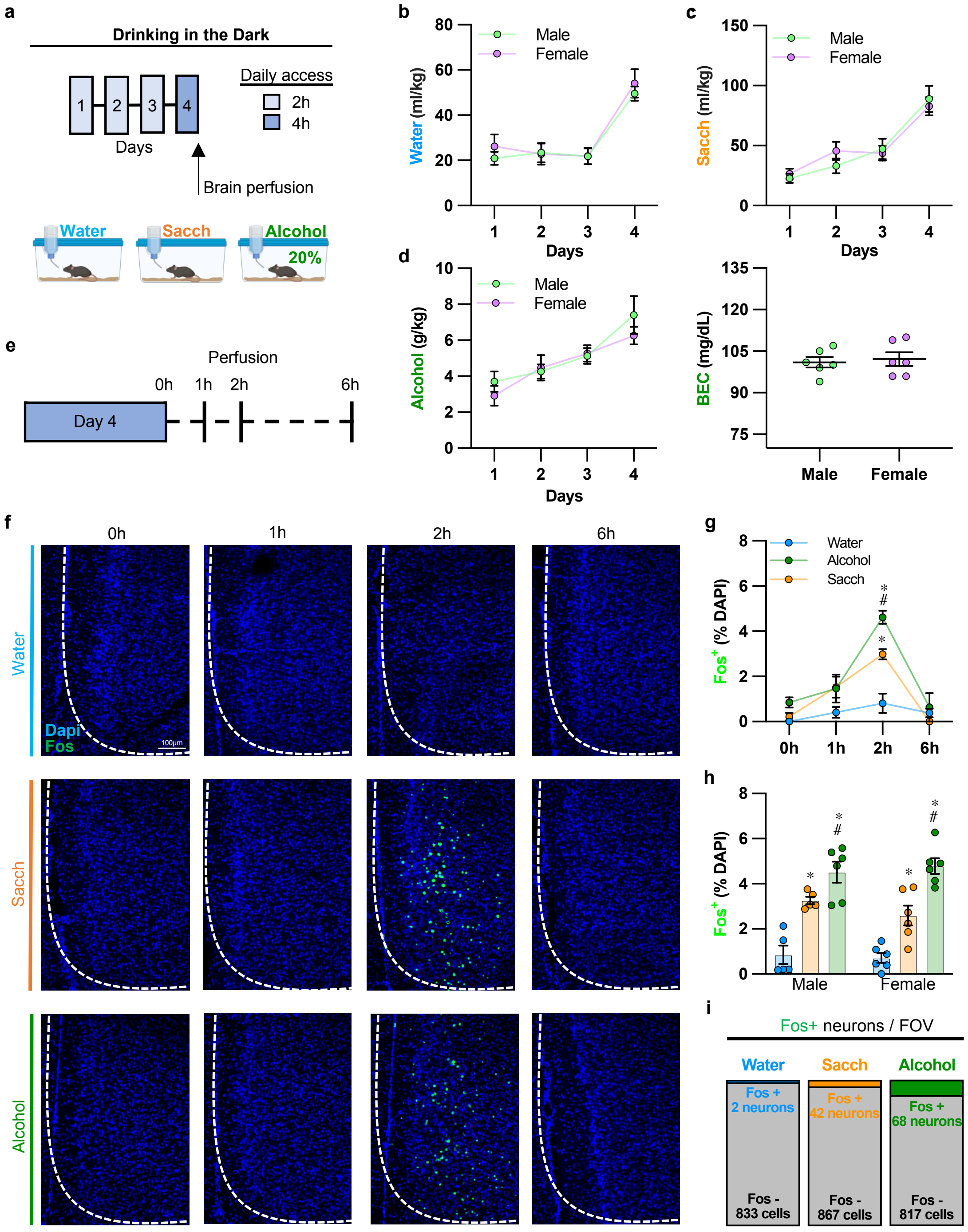


### EXTENDED FIGURE 1

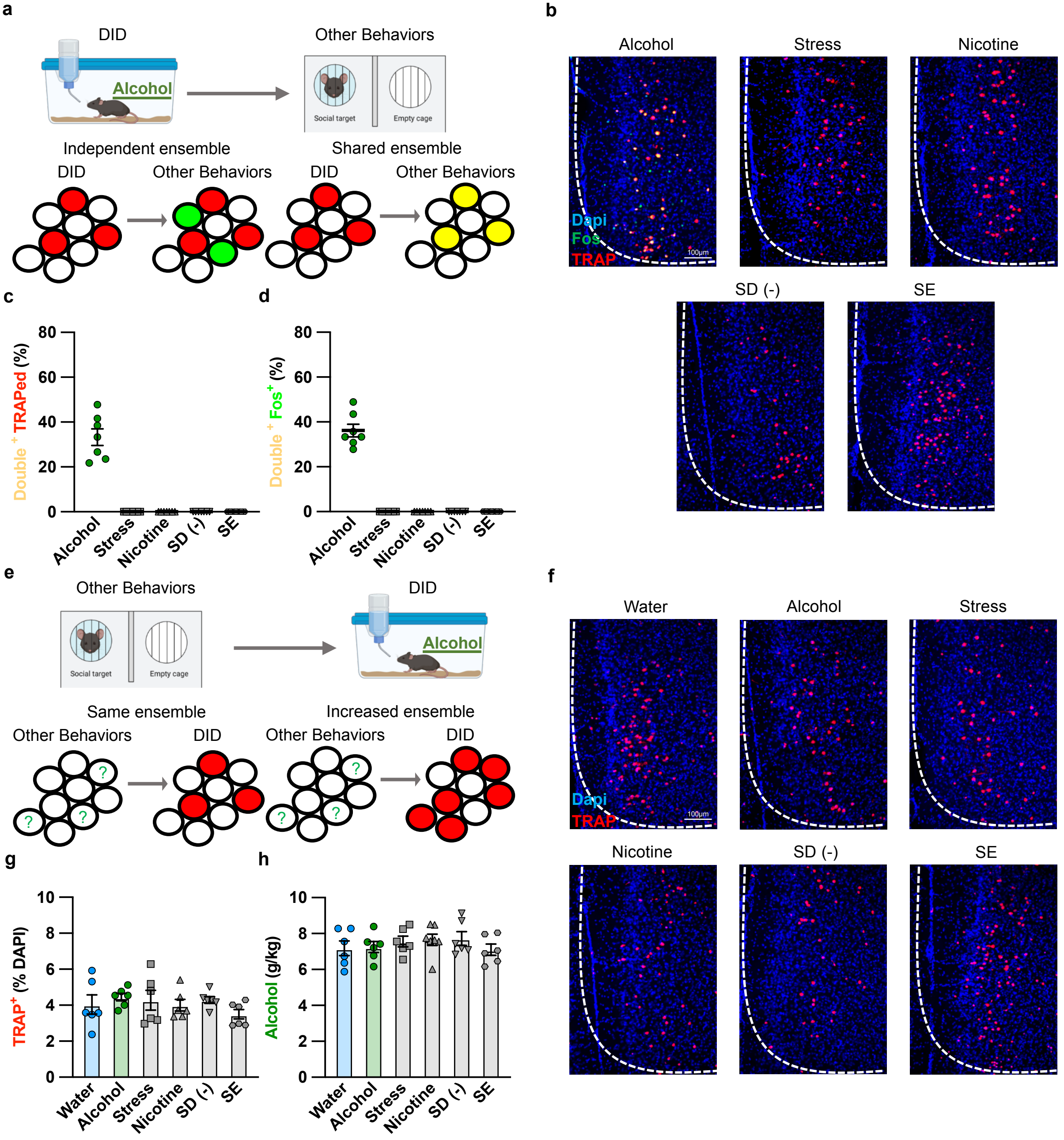


### EXTENDED FIGURE 2

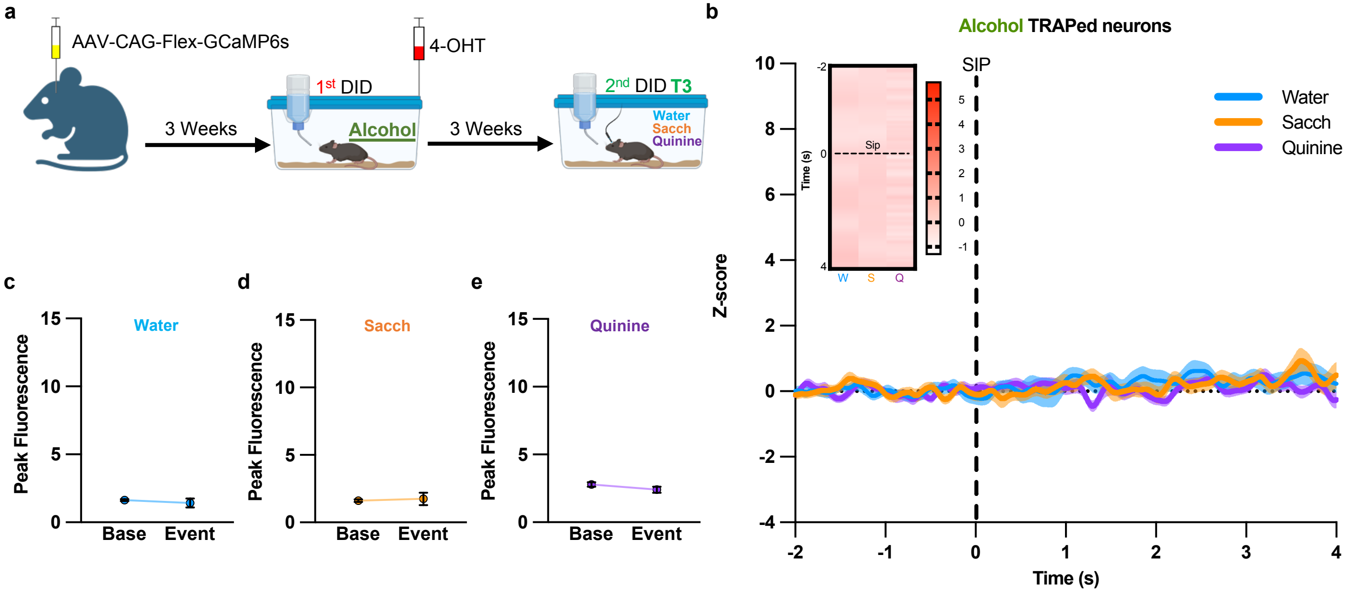


### EXTENDED FIGURE 3

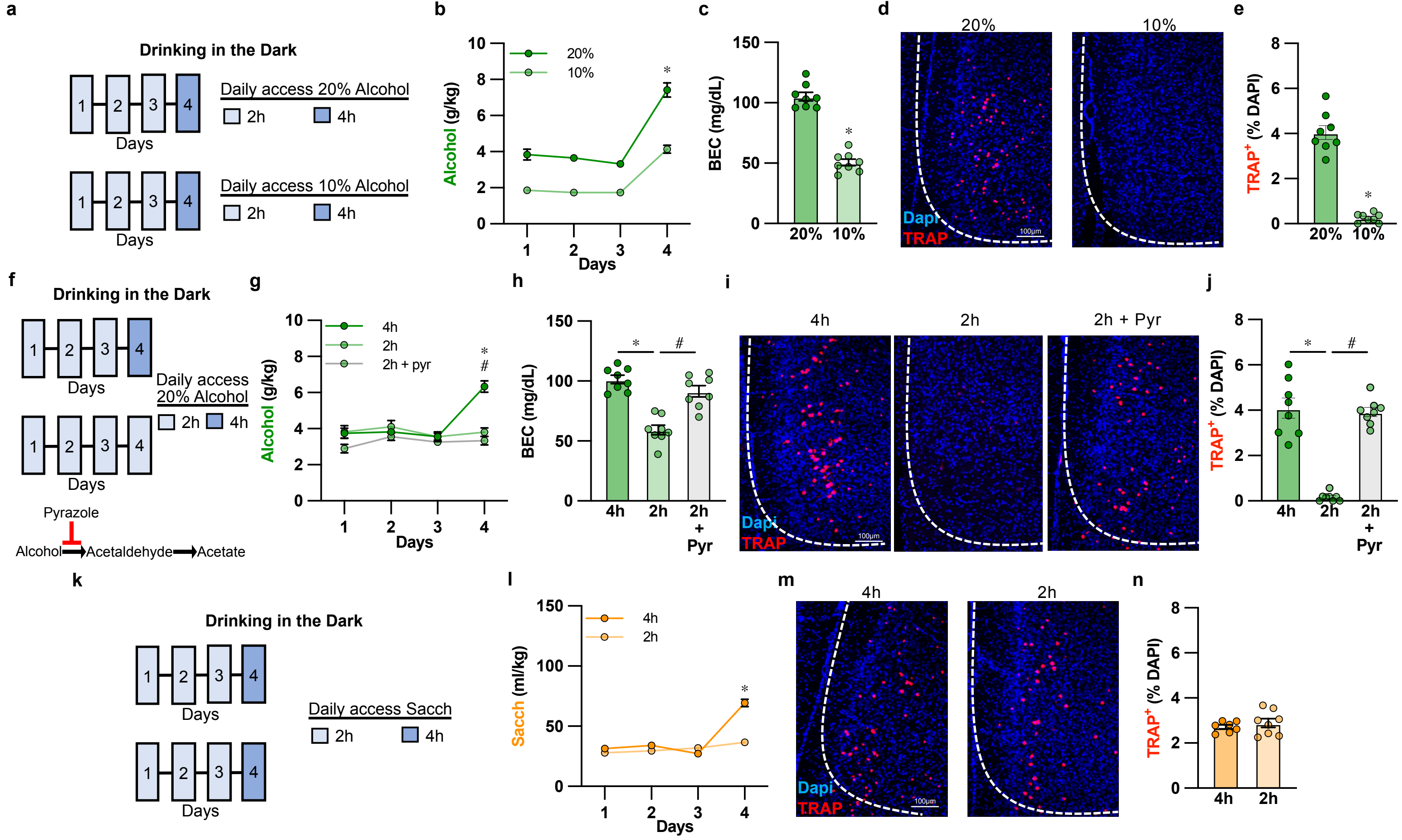


### EXTENDED FIGURE 4

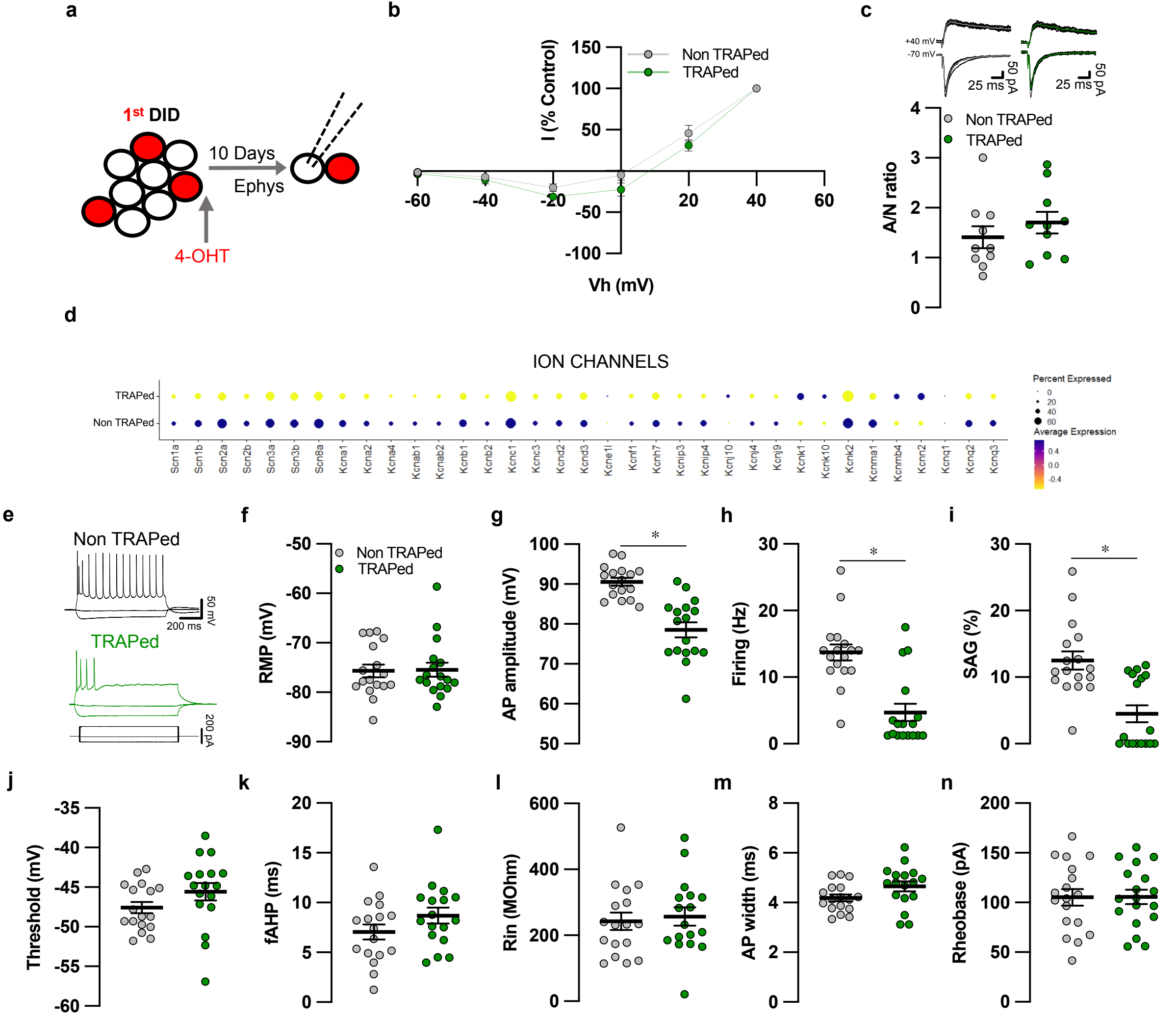


### EXTENDED FIGURE 5

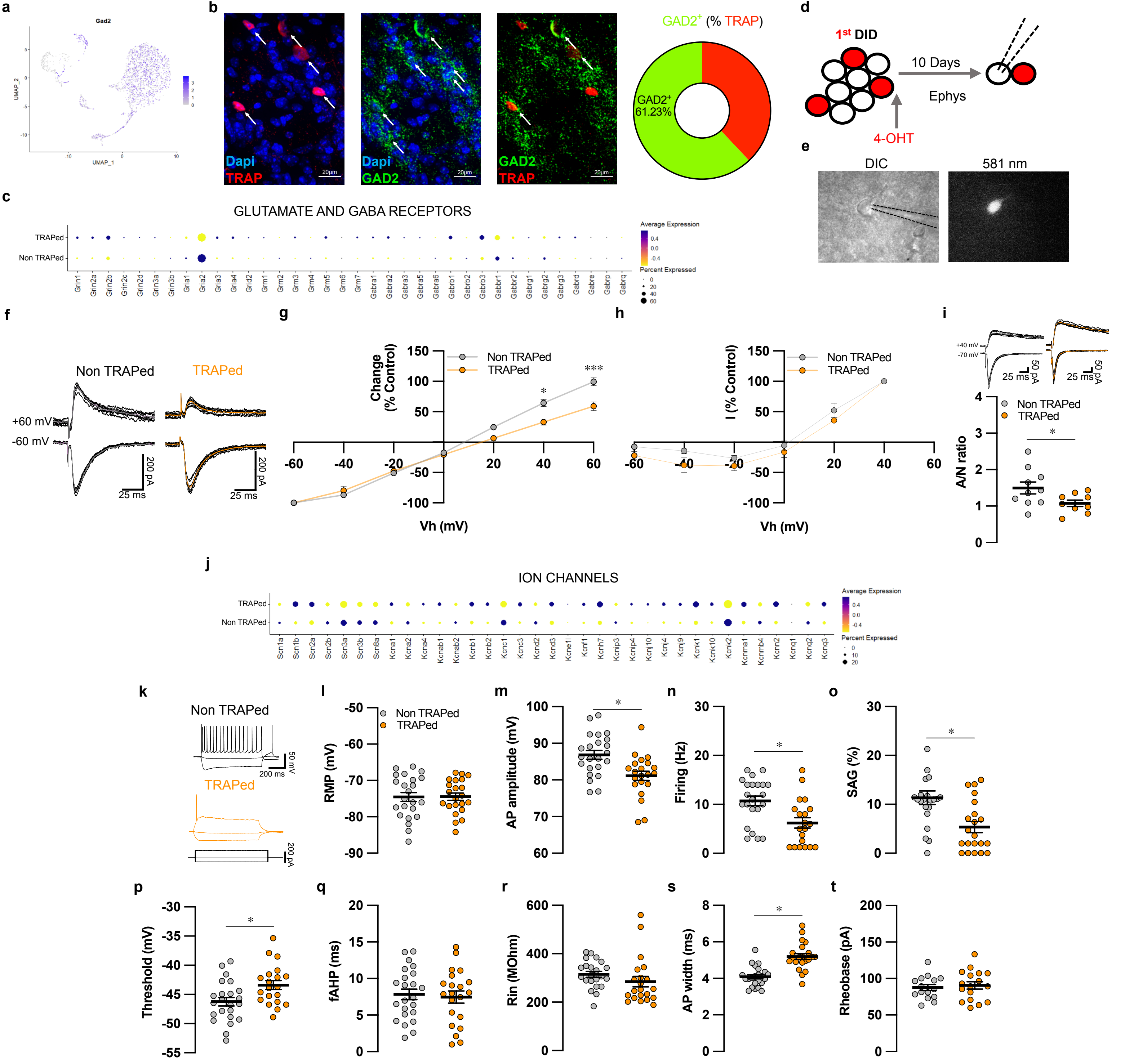


### EXTENDED FIGURE 6

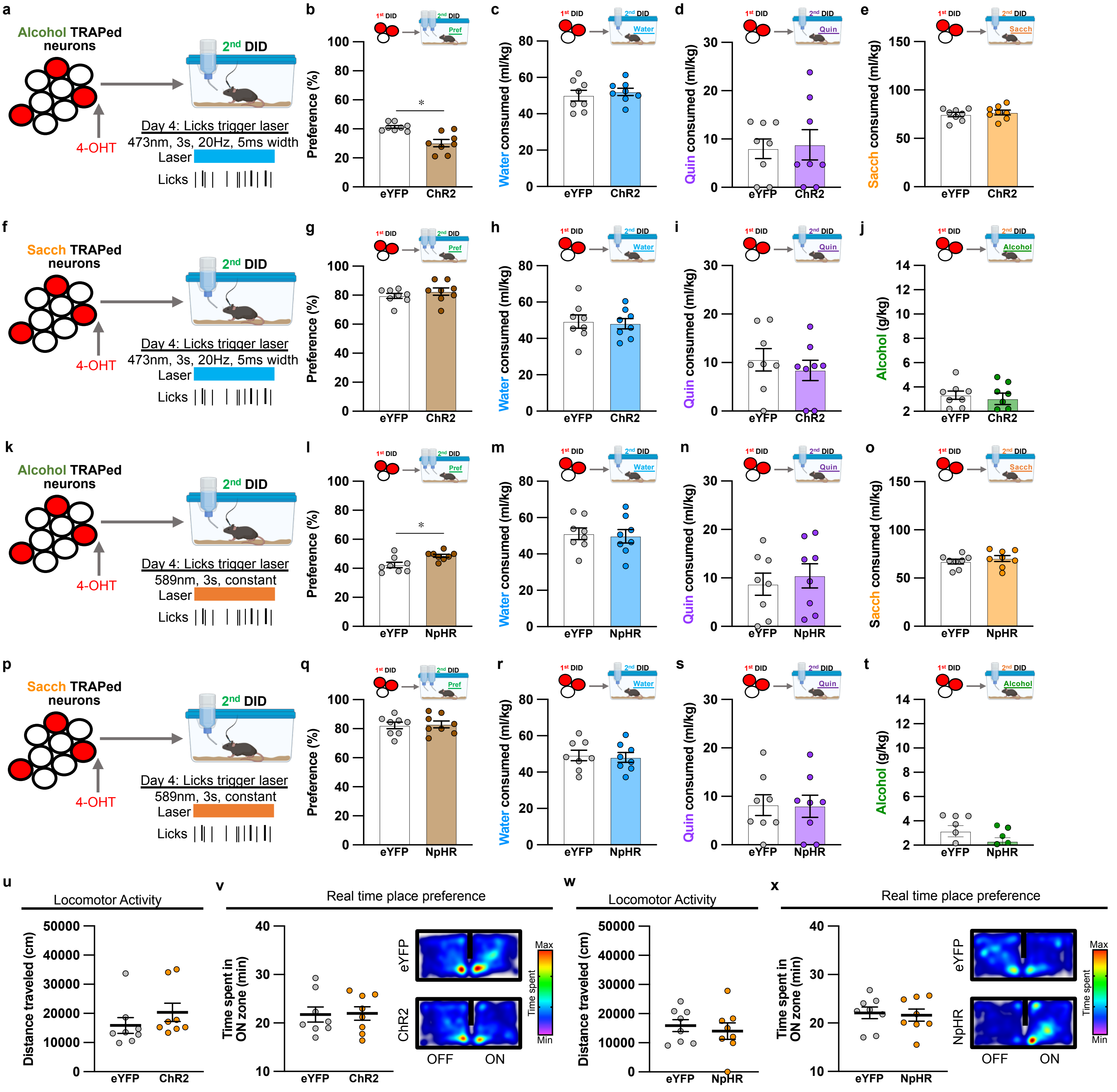


### EXTENDED FIGURE 7

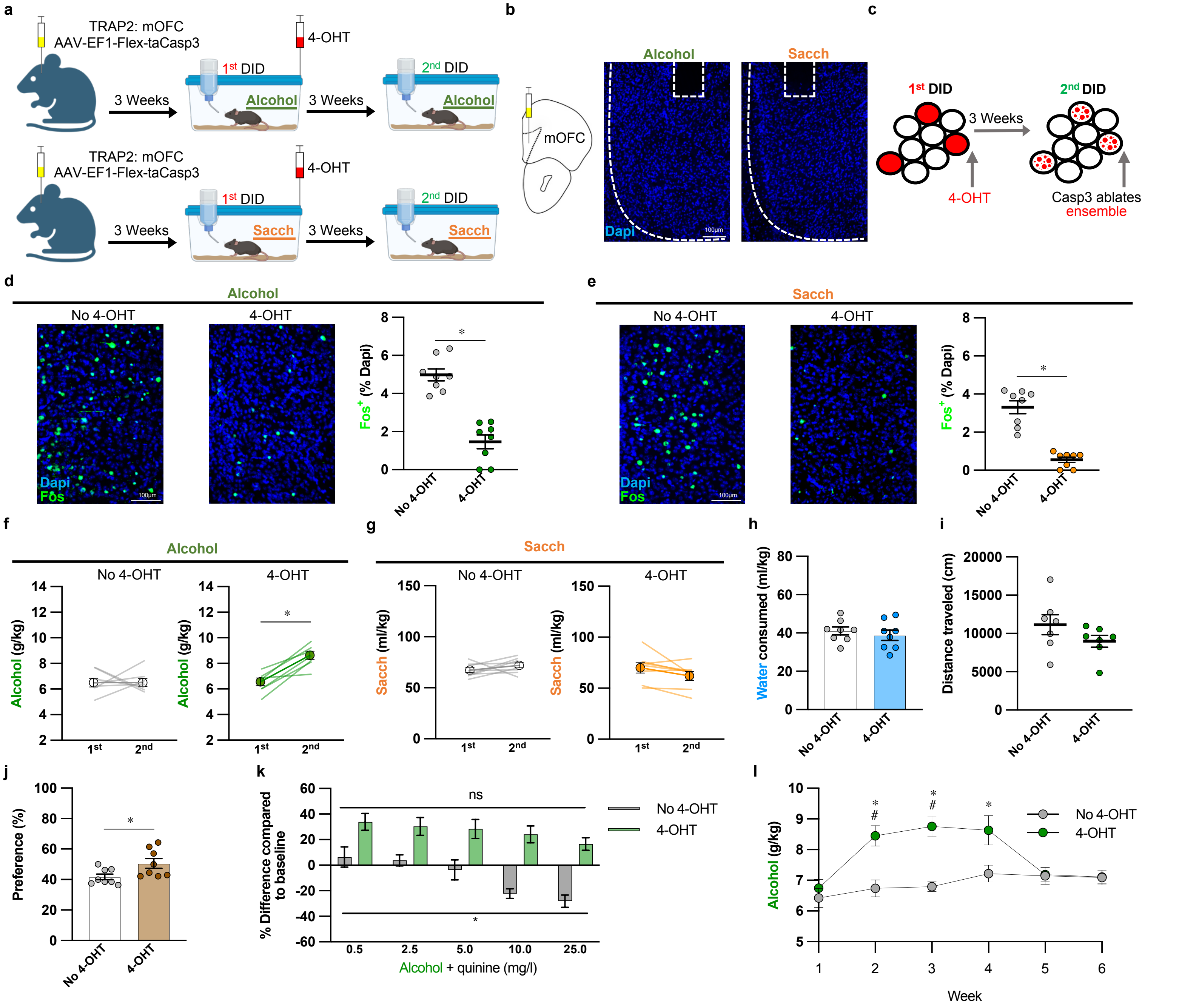


### EXTENDED FIGURE 8

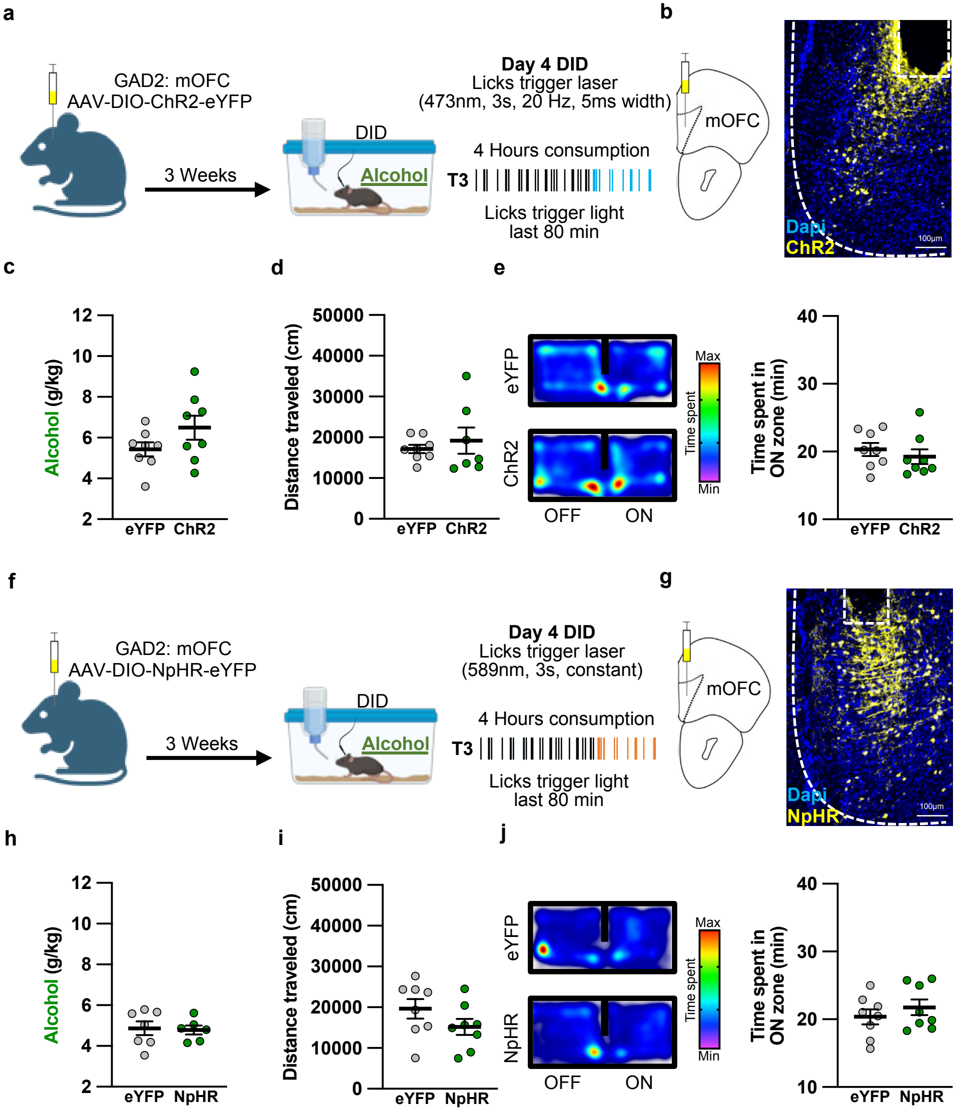


### EXTENDED FIGURE 9

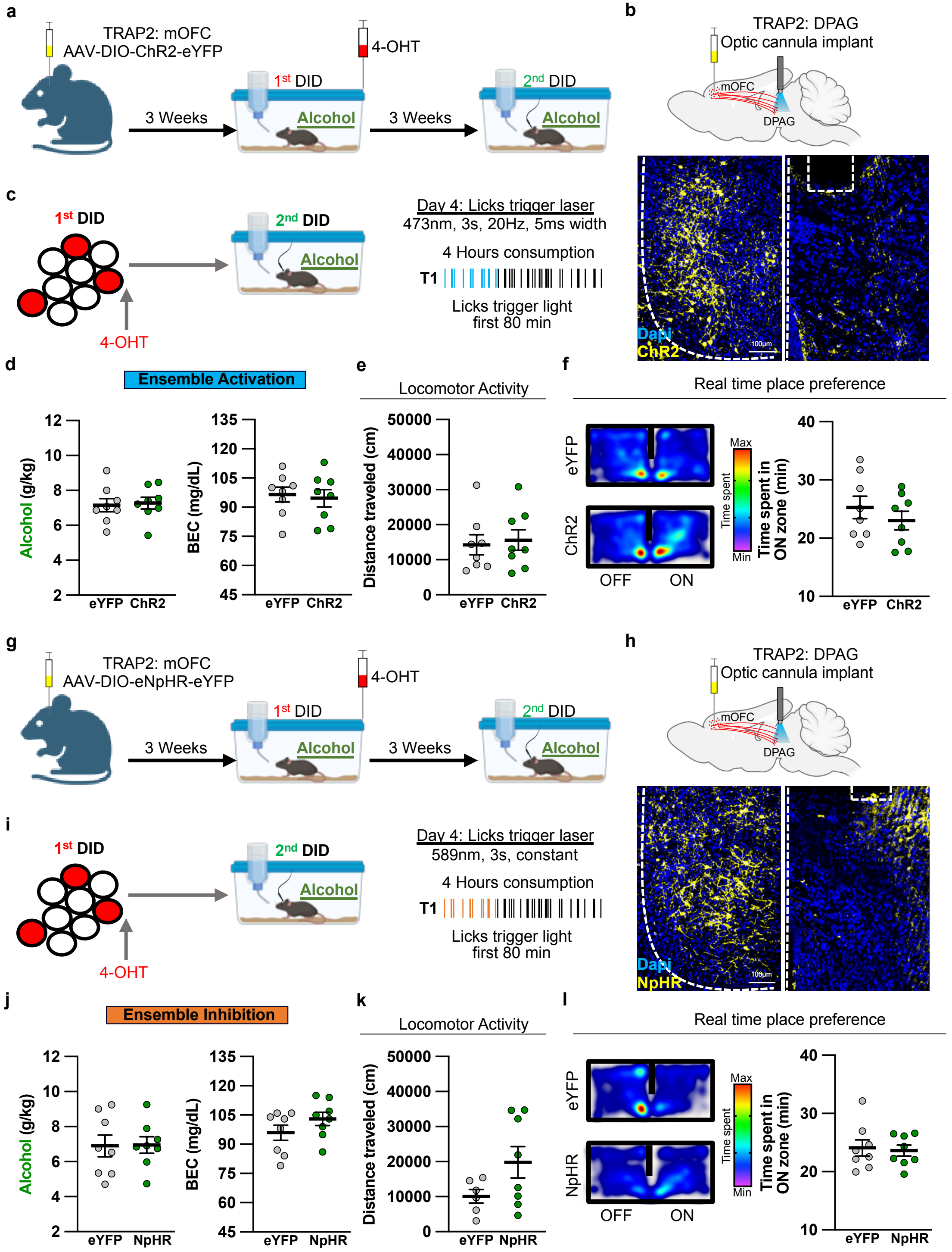


### EXTENDED FIGURE 10

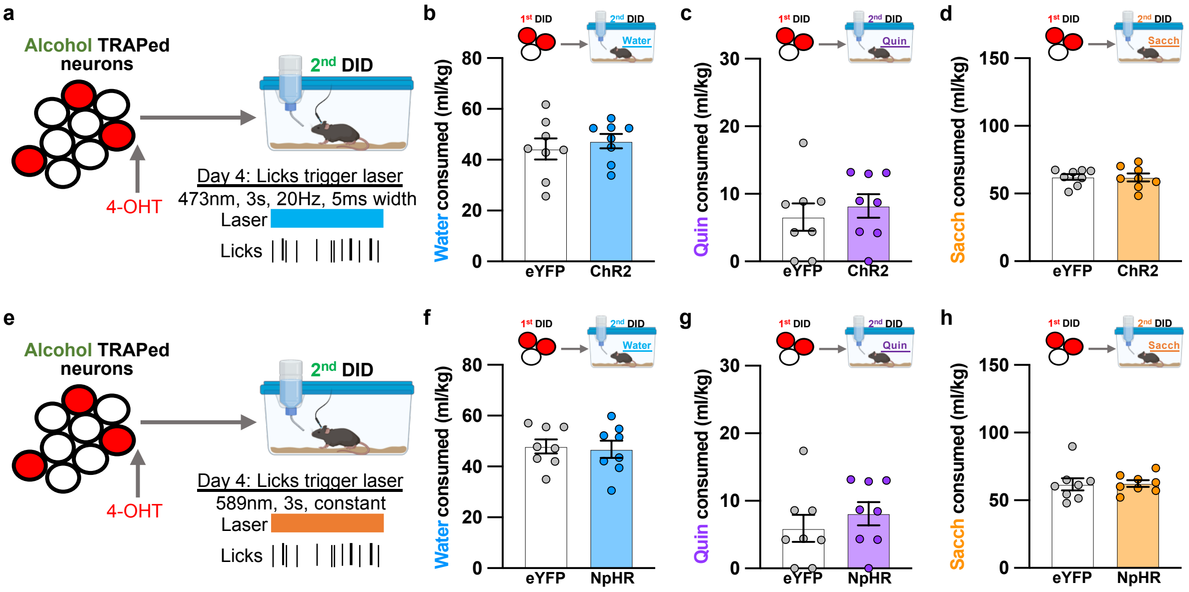


### EXTENDED FIGURE 11
